# Population genetic consequences of the seasonal migrations of birds

**DOI:** 10.1101/2024.06.28.601242

**Authors:** T.M. Pegan, A.A. Kimmitt, B.W. Benz, B.C. Weeks, Y. Aubry, T.M. Burg, J. Hudon, A.W. Jones, J.J. Kirchman, K. Ruegg, B.M. Winger

## Abstract

Differences in life history can cause co-distributed species to display discordant population genetic patterns. In high-latitude animals, evolutionary processes may be especially influenced by long-distance seasonal migration, a widespread adaptation to seasonality. Although migratory movements are intuitively linked to dispersal, their evolutionary genetic consequences remain poorly understood. Using ∼1700 genomes from 35 co-distributed boreal-breeding bird species, we reveal that most long-distance migrants exhibit spatial genetic structure, revealing evolutionary effects of philopatry rather than dispersal. We further demonstrate that migration distance and genetic diversity are strongly positively correlated in our study species. This striking relationship suggests that the adaptive seasonal shifts in biogeography that long-distance migratory species undergo each year lends them enhanced population stability that preserves genetic diversity relative to shorter-distance migrants that winter at higher latitudes. Our results suggest that the major impact of long-distance seasonal migration on population genetic evolution occurs through promotion of demographic stability, rather than facilitation of dispersal.

## Introduction

Co-occurring species provide layers of information about how evolution has shaped the distribution and abundance of genetic diversity across their shared landscape. Spatial genetic patterns can be concordant among co-occurring species, reflecting the extrinsic influence of the shared landscape on evolutionary processes (1). However, intrinsic biological differences between species can also modulate population demographics in ways that influence evolution (2). For example, variation in life history can affect the amount of genetic diversity in populations (3), while differences in dispersal influence the ecological and genetic connectivity of metapopulations (2,4), the dynamics of local genetic adaptation (5), and the formation of new species (6). Comparison of spatial genetic patterns among species on a common landscape therefore provides a rich source of information about how ecological and evolutionary processes are influenced by species’ intrinsic traits (7–9). However, comparative studies involving whole genomes from many co-distributed species have been scarce (10).

To test hypotheses about intrinsic drivers of spatial genetic evolution at a more powerful comparative scale than previously possible, we generated a dataset of 1673 whole genomes sampled from across the geographic ranges of 35 species of birds that co-occur throughout the North American boreal forest (15-67 genomes per species; Supplementary Table 1), sampling that required several decades of field work (Fig. 1). We then used a comparative phylogenetic framework to evaluate intrinsic factors that mediate differences in genetic diversity and patterns of spatial genetic differentiation across the continuous landscape shared by these species.

**Fig. 1.**
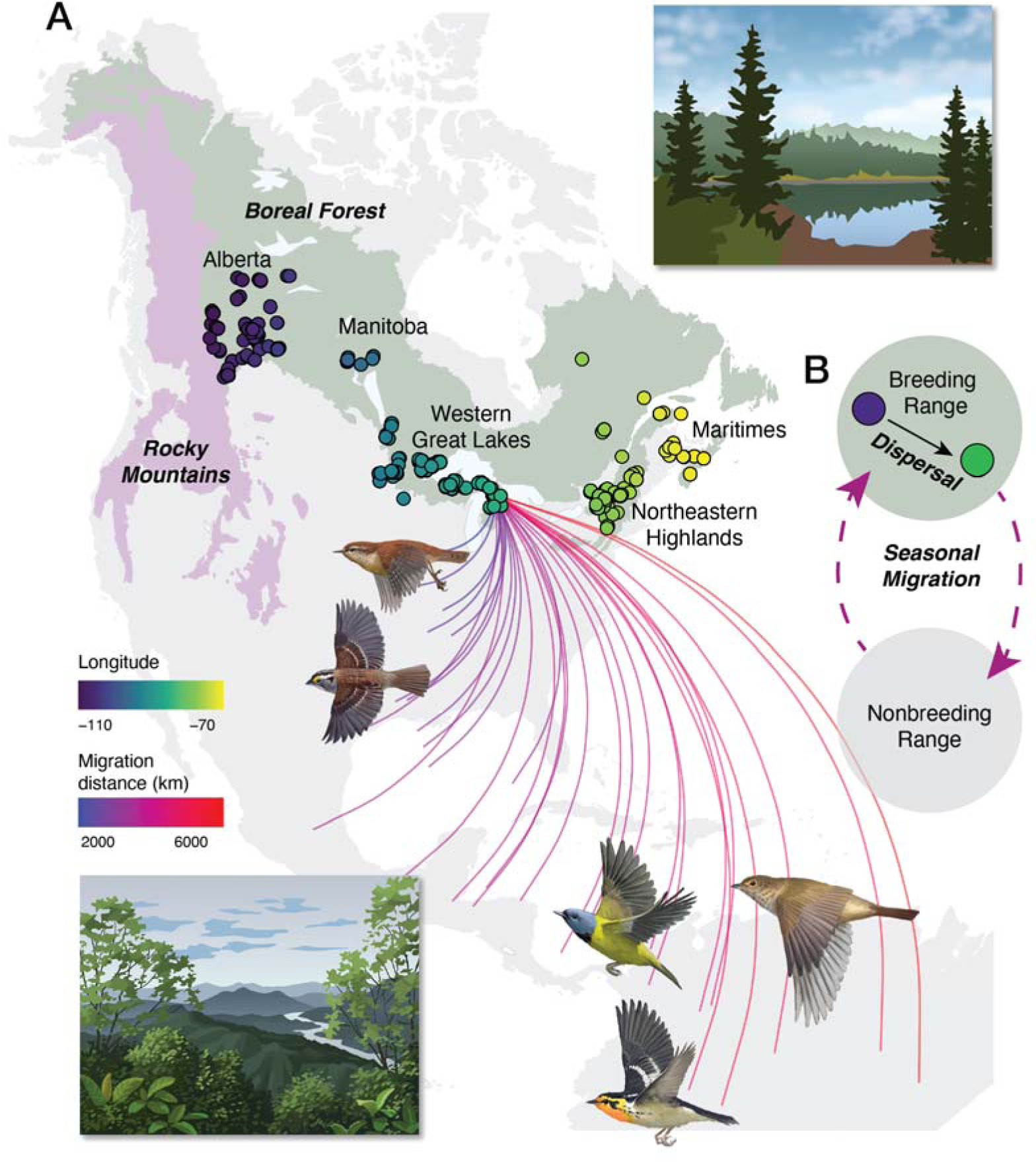
Ecogeographic context of the study. **A**) Study design. The North American boreal ecoregion is shown in green. Points, colored by longitude, show sampling localities along the longitudinal axis of this ecoregion from Alberta, Canada east of the Rocky Mountains to the mainland Maritime provinces of Canada. The 35 species in this study are broadly co-distributed across this region and were sampled evenly across this study area. The nearby Rocky Mountains ecoregion (in pink) contains ecologically similar but often phylogeographically-diverged populations. We did not sample the Rockies nor the extremes of the boreal region in Alaska and Newfoundland, where phylogeographic breaks are known to occur, because of our focus on continuous spatial genetic variation and not all species are found that far east or west. Curved lines depict species-level migration distances of the migratory species in the study, connecting an arbitrary point within their boreal breeding ranges to the centroid of each species’ nonbreeding distribution. **B**) Conceptual diagram of movement behaviors. Dispersal is a movement between breeding sites that can influence gene flow and is distinct from seasonal migration, a round-trip movement between breeding and nonbreeding sites (17). Artwork in panel A by John Megahan. Ecoregion shapefiles from (18).

The ecogeographic setting of our study—the North American boreal forest (Fig. 1)—is particularly well-suited to test hypotheses about how species’ attributes influence their spatial genetic patterns because it represents a geographically vast region that lacks major geographic or environmental barriers, yet is large enough (>3500 km in longitudinal breadth) for spatial genetic patterns to accrue, even in dispersive species (11). Species tend not to show discrete population structure within the boreal ecoregion, whereas strong genetic differentiation between boreal populations and populations in neighboring ecogeographic regions, such as the Rocky Mountains (Fig. 1), is more common (e.g. (12–15)). Within our boreal forest study system, a continuous pattern of isolation by distance is the expected spatial genetic consequence of dispersal limitation, as opposed to deep differentiation driven by barriers to gene flow. This expectation allows us to test whether traits that influence dispersal and population demographic processes affect the presence or strength of isolation by distance patterns across this large biome (6,16).

An important life history trait with potential to shape population genetic evolution in the boreal avian assemblage is seasonal migration, as most bird species breeding at temperate latitudes migrate to lower latitudes after the breeding season to avoid harsh winter conditions (19) (Fig. 1). A species’ migration distance covaries with aspects of biogeography and life history that may influence standing genetic diversity (3,19,20). Additionally, migration is thought to influence dispersal behaviors that can shape spatial patterns of genetic differentiation across the landscape. It is intuitive to assume that long-distance migratory species have high dispersal potential, and there is evidence that migration promotes dispersal in migratory species compared with nonmigratory species (21,22). However, seasonal migration may also promote population divergence and even speciation by increasing behavioral or chronological isolation among populations (reviewed in 23). Further, some studies have noted surprisingly low rates or distances of dispersal in migrants in spite of their high mobility (22,24), and migration appears to restrict geographic range expansion in some contexts (25,26). Such paradoxical restrictions on dispersal in migrants are thought to be a result of high philopatry, i.e., the tendency of migratory birds to return annually to the same breeding location (23,27). Taken together, these observations suggest that migratory birds are characterized by traits that may both promote and suppress gene flow, but it is unclear how these conflicting effects of seasonal migration play out to influence spatial genetic differentiation over evolutionary time.

To shed new light on the evolutionary consequences of seasonal migration, we leverage the considerable variation among boreal-breeding bird species in their nonbreeding ranges and thus their seasonal migration distances (Fig. 1). We first test the hypothesis that migratory boreal species have lower spatial genetic structure than the nonmigratory species that are resident year-round in the boreal region, due to their higher mobility. Further, using only migratory species, we test the hypothesis that spatial structure scales negatively with migration distance such that the longest-migrating species experience the highest gene flow within their breeding range and therefore demonstrate the lowest spatial structure.

In addition to evaluating the effects of seasonal migration on spatial genetic patterns, we evaluated two other species attributes that may influence spatial genetic differentiation: 1) body size, which is positively correlated with dispersal distance in birds (21); and 2) species’ associations with early successional boreal habitat (as opposed to mature boreal forest), because habitat ephemerality is associated with lower levels of population genetic structure (28). We also included an estimate of population-wide genetic diversity (θ_π_) as a third covariate to test whether genetic diversity predicts spatial genetic structure across species. A positive correlation may occur because low rates of dispersal between metapopulations can promote both genetic diversity and spatial genetic differentiation (29). Moreover, in post-glacial landscapes, population expansion and invasion dynamics can affect both genetic diversity and the formation of spatial genetic structure in complex and interacting ways (30). Such effects may be especially relevant in boreal populations because their breeding ranges were repeatedly shifted during glacial-interglacial cycles of the Pleistocene and they have inhabited their contemporary spatial context for no more than 20 thousand years, following the end of the Last Glacial Maximum (31).

Finally, the variation in migration distance among our study species coupled with the absence of strong geographical barriers provides a unique opportunity to test how migration, a key life history adaptation to seasonally fluctuating environments, influences standing genetic diversity in populations occupying the highly seasonal boreal latitudes. Genetic diversity is an important parameter that reflects the size and demographic dynamics of populations over their evolutionary history (32) and carries implications for future population persistence (33). Prior work suggests that life history traits, including migratory behavior, influence evolutionary processes that shape genetic diversity (3,34,35), but the mechanisms by which life history parameters influence genetic diversity remain poorly understood (36,37). As the long-distance migrants in our study have been shown to have a slower life history than their codistributed short-distance migrants (19), our study design provides a detailed test of the relationship between life history and genetic diversity among co-distributed species sampled in a consistent manner from a common landscape.

## Results and Discussion

### Migration and spatial genetic structure

Our analyses reveal that boreal bird species vary considerably in spatial population genetic patterns and genetic diversity despite their common landscape and broadly similar histories in the region (38) (Fig. 2, Fig. S1-S35, table S1), presenting the opportunity to test whether interspecific variation in these patterns can be explained by species’ attributes. To test predictors of spatial genetic structure, we used a multilevel modeling approach (‘HMSC;’ 39) that estimates the relationship between predictor variables and the dependent variable within species while jointly estimating the effects of species’ attributes on interspecific variation and accounting for phylogenetic relatedness among species. Specifically, we used HMSC to model the relationship between pairwise genetic covariance and pairwise geographic distance in all conspecific sample pairs, generating a slope of isolation by distance *(β_IBD_*) for each species (Fig. 3), while simultaneously modeling the effects of all four species attributes—migration, mass, habitat association, and genetic diversity—on isolation by distance (Fig. 3, γ coefficients). Under a model of isolation by distance, genetic covariance values are expected to be positive in geographically close pairs of samples and negative in geographically distant pairs that are more distantly related (16,40). As such, *β_IBD_* coefficients, which describe the slope of the IBD curve, are negative in species demonstrating isolation by distance, and species attributes with negative γ parameters are those associated with spatial partitioning of genetic structure.

**Fig. 2.**
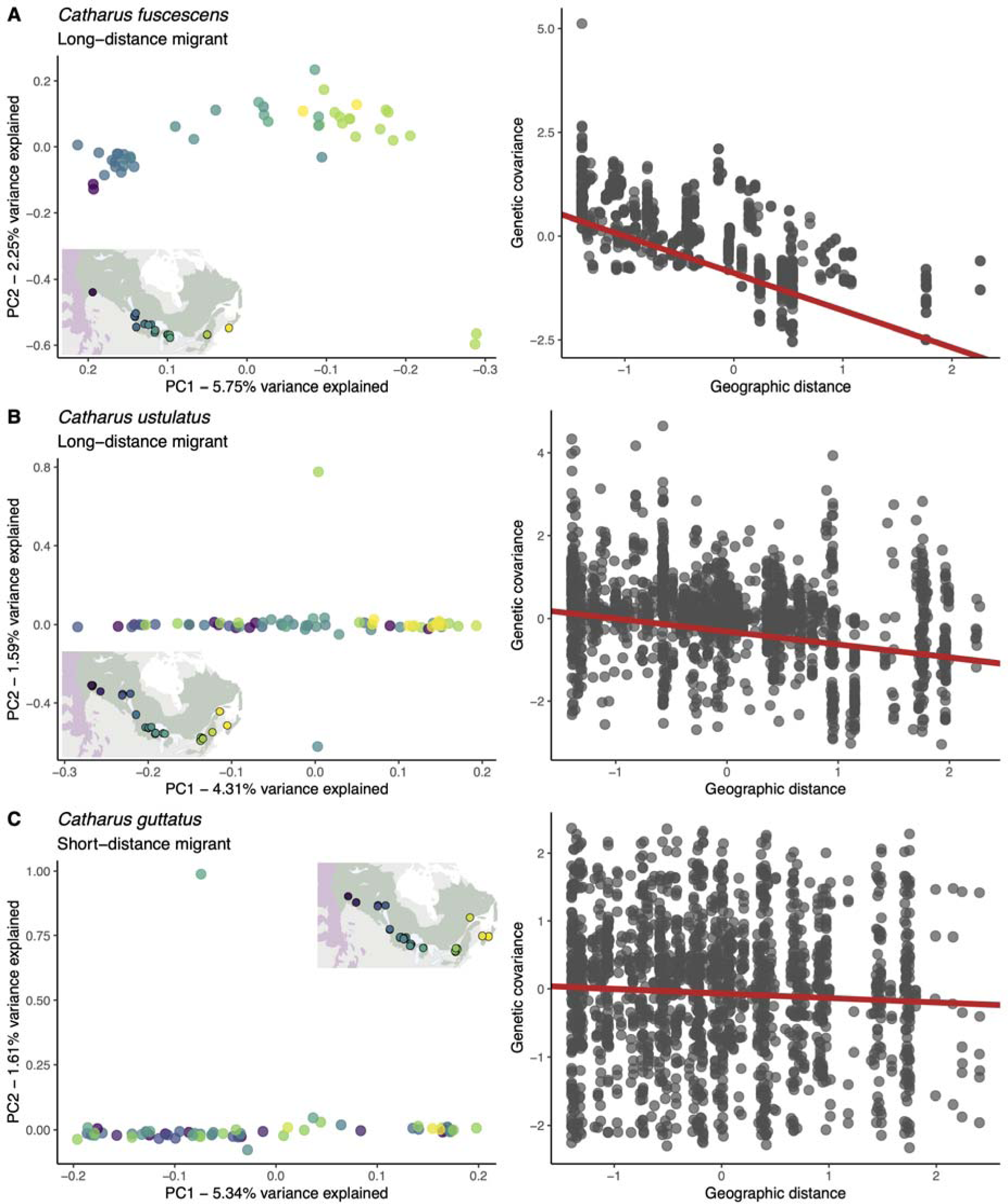
Interspecific variation in spatial genetic patterns. PCA and *β_IBD_* plots from three example species in the genus *Catharus* demonstrate the variety of spatial genetic patterns present in boreal populations of the 35 study species (Fig. S1-S35). The long-distance migrant *C. fuscescens* (**A**) shows a clear spatial genetic pattern of isolation by distance, whereas *C. ustulatus* (**B**) shows subtler isolation by distance and the shorter-distance migrant *C. guttatus* (**C**) shows no spatial structure. Point colors in PCA plots correspond to longitudinal locations of samples, as shown in the inset of each panel. Each PCA is accompanied by a scatterplot showing the relationship between genetic covariance and geographic distance, with the slope and intercept of *β_IBD_* for that species (*Materials and Methods*; Fig. 3).

**Fig. 3.**
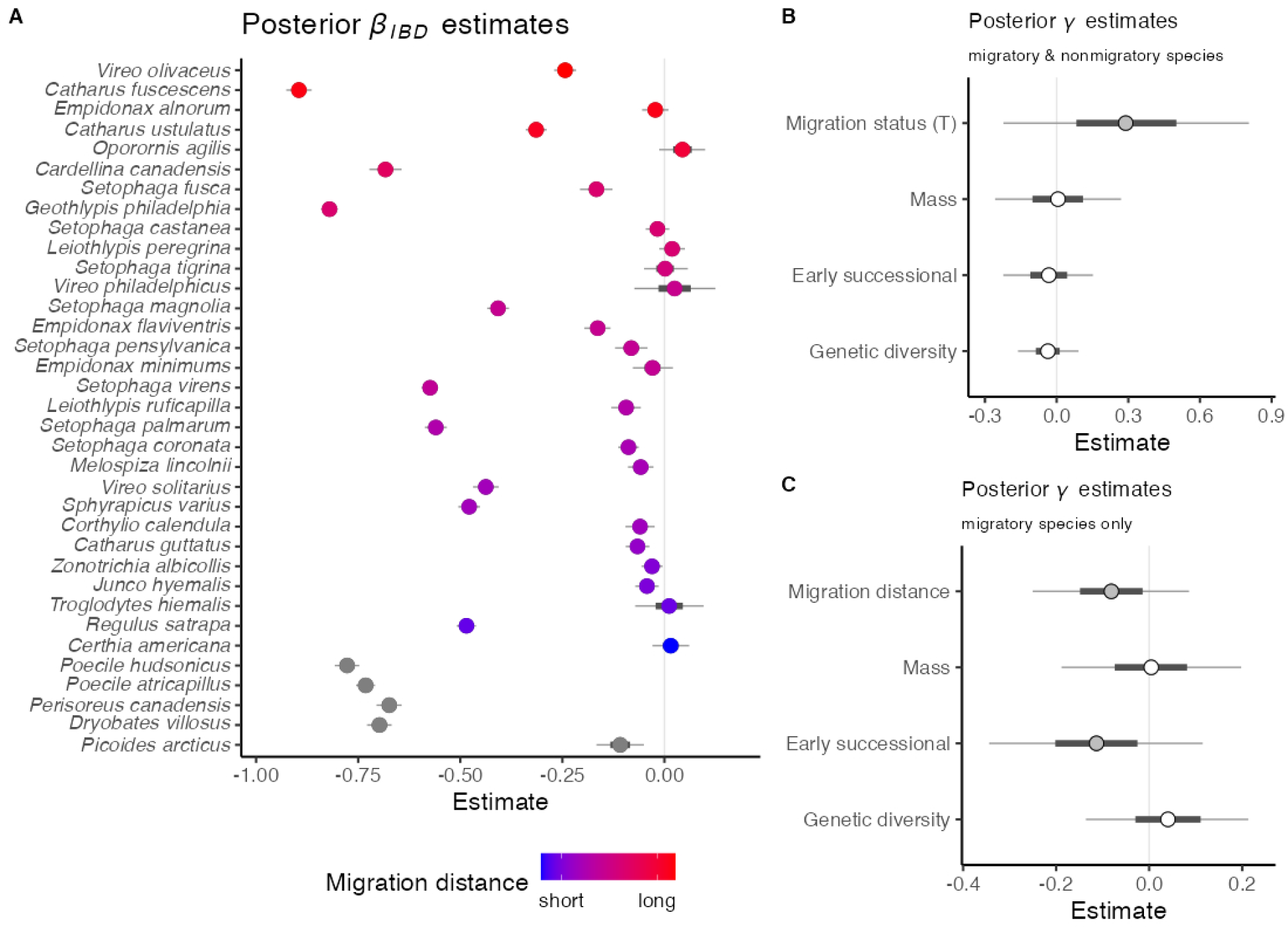
Isolation by distance (*β*_IBD_) and effects of species attributes on *β*_IBD_. Posterior estimates of *β_IBD_* (**A**) and γ (**B, C**) and their credible intervals from HMSC models. Thick lines represent 50% credible intervals and thin lines represent 90% credible intervals. A) Species-specific slopes of isolation by distance (*β*_IBD_) generated from the HMSC hierarchical mixed models. We created one model where migration was coded as a binary trait, and another including only the 30 migratory species with migration distance as a continuous trait. *β_IBD_* estimates were highly similar across models, so we show results only from the full 35-species model, with points colored by migratory and migration distance (nonmigratory = gray). Species are arranged on the y axis in order of migration distance. Most species show negative *β_IBD_* coefficients, reflecting the negative relationship between genetic covariance and geographic distance expected under a model of isolation by distance. Other species show *β_IBD_* coefficients close to 0, reflecting a weak or absent relationship between the spatial distribution of samples and their genetic covariance. Posterior γ estimates from HMSC models (**B, C**) reflect the influence of traits on community-level variation in *β_IBD._* In each case, a negative value of γ indicates that a trait promotes spatial structure (i.e. results in a more negative *β_IBD_*), while a positive value of γ indicates that a trait reduces spatial structure. The central point is colored by the position of 0 relative to the estimate’s credible intervals, where 0 is within the 50% interval for white points and outside of it for gray points.

We tested our two hypotheses about the effects of seasonal migration on spatial genetic structure using two models. The first model contained all 35 species in the study and included a binary γ variable indicating migratory status (migratory vs nonmigratory). We used the second model to test whether effects of migration scale with migration distance (as a continuous variable); this model contained only migratory species (*n* = 30). We included the other three attributes (mean mass, genetic diversity, and a binary variable indicating association with early successional habitat; Fig S36) as γ variables in both models.

We found that nonmigratory species tend to display clearer spatial structure within the boreal region than migratory species, suggesting that migratory species experience higher rates of gene flow. Four out of the five nonmigratory species in our study show clear spatial patterns in PCA plots, whereas many migratory species lack an evident geographic pattern (e.g., Fig. 2C, Fig. S4-S6, S25-S30, S33-S34). HMSC models also provide modest support for an effect of migratory status on spatial structure, where migrants are predicted to show weaker (less negative) *β_IBD_* (mean posterior γ = 0.29, posterior support = 0.82; Fig. 3). Our conclusions about nonmigratory behavior are necessarily based on data from few nonmigratory species, as most boreal bird species migrate. However, these conclusions are consistent with other studies that used mark-recapture data to demonstrate that nonmigratory birds typically undergo reduced dispersal compared to migratory birds (21,22).

Although our results suggest that migratory birds experience more extensive gene flow than nonmigratory birds, we found a contrasting effect of migration distance among the migratory species. In the model including only migratory species (*n* = 30), long-distance migration was not associated with increased gene flow. Rather, the γ coefficient for an effect of migration distance on *β_IBD_* was negative (γ = −0.081, posterior support = 0.8, Fig. 3), suggesting somewhat higher spatial genetic differentiation in long-distance migrants relative to short-distance migrants. Although posterior support for this modeled relationship was modest, our results make clear that some extremely long-distance migratory species in our study exhibit notable spatial structure or isolation by distance (e.g. Fig. 2C, S23, S24, S32, S35). These patterns may reflect limited gene flow across the boreal ecoregion, a remarkable pattern to discover in species that annually migrate distances roughly 1.5 to 2 times the entire breadth of the boreal breeding ranges where gene flow could occur.

Spatial genetic structure in long-distance migratory species may reflect a counterintuitive relationship between seasonal migration distance and philopatry (17). While long-distance movement tends to be the most obvious and remarked-upon characteristic of seasonal migration, philopatry is equally fundamental—it is what distinguishes migration from other movement behaviors such as nomadism and dispersal (27). When time in the breeding range is most limited, there is pressure to establish a breeding attempt quickly (41) and there are clear advantages of returning to a familiar area, rather than exploring elsewhere (42,43). This effect may be strongest in long-distance migrants, which spend the least time in their breeding regions (19) (Fig. S37).

Spatial structure in long-distance migratory species may therefore reflect an evolutionary consequence of the remarkable navigational precision that long-distance migrants employ to travel around the globe only to return to the exact same breeding sites year after year (e.g., 44,45).

### Other predictors of spatial structure

Aside from migration, the evidence for effects of other species attributes on population genetic patterns in the boreal avian assemblage was less clear. In our HMSC models, posterior support values for effects of genetic diversity (a proxy for *N_e_*) on *β_IBD_* were equal to or less than 0.7 in all cases, which means that our models do not support a correlation between genetic diversity and spatial genetic structure. We found similarly low support for hypothesized effects of body mass on spatial structure. We predicted that species associated with habitat in early stages of succession would show reduced spatial genetic structure because early successional habitats are spatially ephemeral compared to mature habitats, as studies in tropical bird species suggest that habitat ephemerality promotes gene flow and reduces signatures of spatial genetic structure (28). However, our HMSC model using only migratory species demonstrated the opposite pattern, wherein early successional habitat was associated with stronger *β_IBD_* (i.e. stronger spatial structure; γ = −0.11, posterior support = 0.81, Fig. 3), and posterior support for this effect was substantially lower in the model with all species (γ = −0.033, posterior support = 0.61, Fig. 3).

Overall, our HMSC models explained only a small proportion of interspecific variation in *β_IBD_* (proportion of variance explained by effects of γ coefficients was 0.010 in the model with all species, and 0.03 in the model with migratory species only), but the models also identified strong phylogenetic signal in residual (unexplained) variation. The mean posterior ρ coefficients were 0.86 ± 0.09 and 0.86 ± 0.10 for the 35-species and 30-species models respectively (where ρ can range from 0, indicating no phylogenetic signal, to 1, indicating very strong phylogenetic signal (46)). The presence of strong phylogenetic signal in residual interspecific variation in isolation by distance raises new questions about how such signal could arise. Whereas inherited phenotypic traits are passed directly from ancestors to descendants, creating a strong expectation of phylogenetic signal in many phenotypes (47), spatial genetic patterns arise emergently through a fundamentally different process of interaction between individuals and the landscape. That boreal species demonstrate phylogenetic signal in their residual spatial genetic variation suggests that unidentified, phylogenetically conserved traits may have a strong influence on how these patterns form (46).

A broad conclusion of our analyses is therefore that much of the interspecific variation in spatial structure among boreal bird species may be driven by species’ intrinsic attributes, rather than extrinsic processes that are random with respect to phylogeny. Although coarse-scale attributes such as general locomotory capacity predict spatial genetic patterns at broad phylogenetic scales (48), our results show that their explanatory power is lost within assemblages of closely-related species, highlighting the difficulty of identifying attributes that play the most important roles in spatial evolution without more refined ecological and natural history information. Specifically, among our study species, we posit that much of the unexplained variation in spatial structure may be related to other aspects of dispersal behavior that are difficult to measure and thus could not be confidently included in the study as covariates. For example, another axis of dispersal behavior in boreal birds is the extent to which these species follow resource variation across their breeding grounds when choosing nesting sites instead of exhibiting philopatry. Indeed, it is noteworthy that the three species of warblers in our study whose population ecology is closely tied to spruce budworm outbreaks (*Leiothlypis peregrina, Setophaga castanea, Setophaga tigrina*; 49) show no discernible geographic structure in PCA (Fig. S25, S28, S34). This result raises the possibility that species that follow prey with boom-bust population cycles may undergo long-distance dispersal in response to food availability, resulting in high gene flow and lack of spatial genetic structure.

### Migration and genetic diversity

Our results also revealed an unexpectedly strong positive correlation between genetic diversity and migration distance (Fig. 4). We found that nonmigratory species tend to show the least standing genetic diversity in the boreal avian assemblage, implying that they have the smallest effective population sizes (*N_e_;* 36), and that genetic diversity (and *N_e_*) increases with migration distance (Phylogenetic Generalized Least Squares effect size = 0.80, std error = 0.10, p < 0.00001, λ = 0.15). García-Berro *et al.* (34) previously showed that migration promotes genetic diversity based on comparisons between migratory and nonmigratory butterflies, and our study demonstrates further that the effect scales with migration distance within temperate-breeding bird species (Fig.4).

**Fig. 4.**
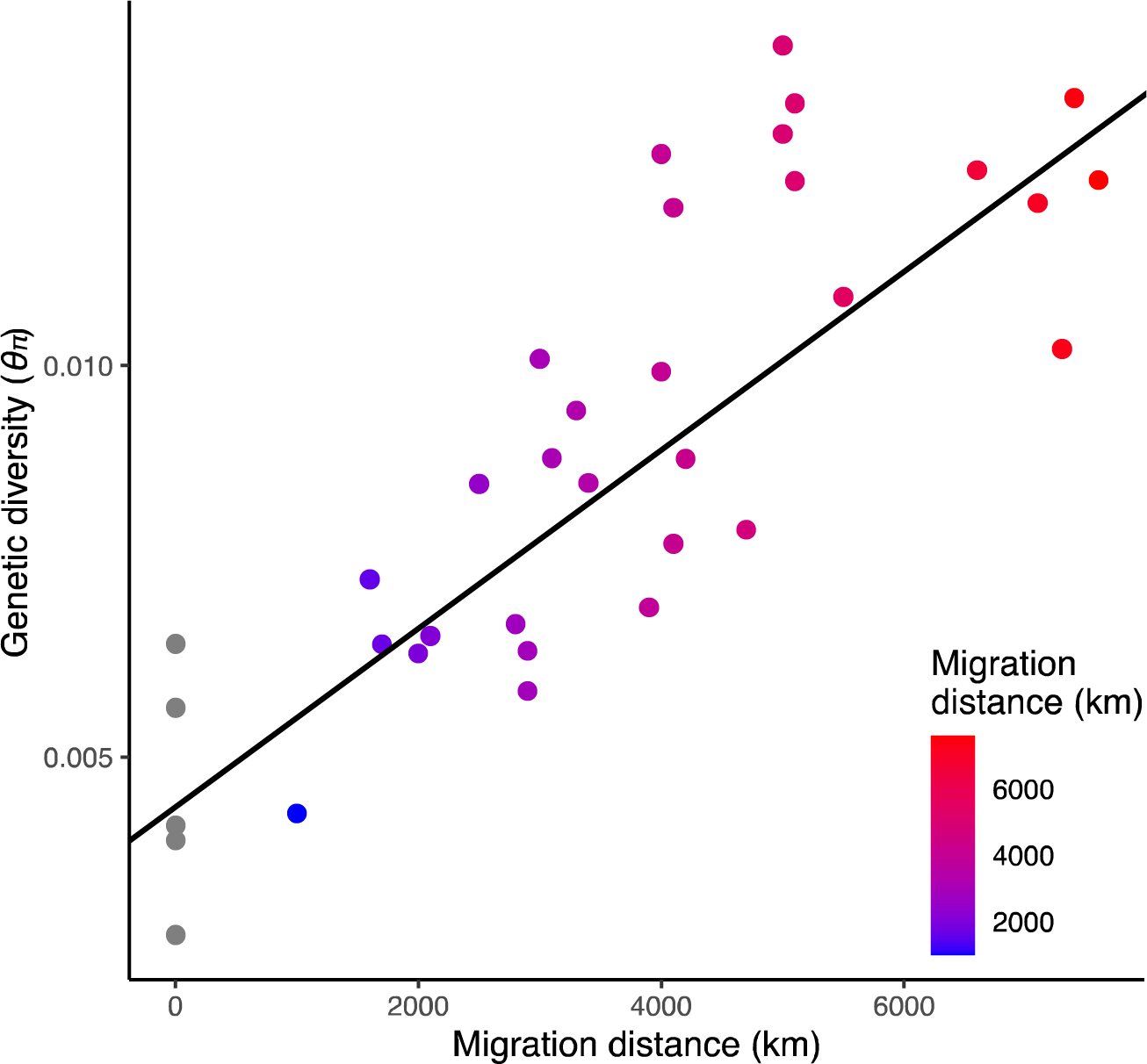
Genetic diversity (θ_π_) correlates with migration distance. Each point represents one species and points are colored by migration distance, as in Fig. 1 and Fig. 3, where non-migrants are shown in gray. The intercept and slope of the line were estimated with a phylogenetic generalized least squares (PGLS) model (*Materials and Methods*).

Why do long-distance boreal migrants have higher genetic diversity? Long-distance migrants in this system also have slower life history (higher annual survival, lower annual fecundity) than short-distance migrants (19). Thus, the genetic diversity pattern we document (Fig. 4) contrasts with results from comparisons across animal taxa at much broader phylogenetic scales, which show that species with slow life history exhibit low genetic diversity (3). However, the mechanisms underlying a relationship between life history and genetic diversity are complex and poorly understood (36,37). Genetic diversity is also thought to be influenced by effects of the environment on demography, including population bottlenecks (32) and unstable metapopulation dynamics (50). Given the overlap in the breeding ranges of our study species, we propose that interspecific variation in genetic diversity in boreal birds is strongly influenced by demographic consequences of the non-breeding environment.

Although recent work suggests that many boreal bird species did not experience recent severe bottlenecks at the Last Glacial Maximum (38), genetic diversity can also be eroded by frequent extinction-recolonization events in metapopulations, which create successive local bottlenecks and reduce genetic variation (51). In particular, boreal nonmigratory species and short-distance migrants are exposed to challenging temperate-zone winter conditions during the nonbreeding season, which are known to limit population sizes (52,53). By contrast, long-distance migrants in our system evade temperate winter by migrating to warm and resource-rich tropical regions, where they experience higher apparent annual adult survival than short-distance boreal-breeding migrants that winter at high latitudes (19) (Fig. 1). Therefore, greater population stability on tropical compared to temperate nonbreeding grounds may buffer long-distance migrants from the frequent loss of genetic diversity. The relationship between nonbreeding latitude and genetic diversity among sympatrically breeding boreal bird species parallels a global relationship between latitude, genetic diversity, life history and population demographic stability that is present in many taxa (20), highlighting the importance of nonbreeding ranges for migratory bird population demographics across ecological and evolutionary timescales. Moreover, our results suggest that in assemblages of migratory birds, interspecific variation in genetic diversity is strongly influenced by the demographic effects of seasonal biogeography.

### Conclusions

Our results carry important implications for how interspecific variation in seasonal migration—a widespread adaptation to seasonal environments—interacts with evolutionary processes. We show that even long-distance migrants can exhibit sufficiently high philopatry to result in spatial segregation of breeding populations across a continuous landscape. Moreover, our comparative framework reveals a previously unknown positive correlation between migration distance and genetic diversity, suggesting that migration to tropical forests each year has allowed long-distance migrants to experience greater long-term population stability than species whose annual cycles take place entirely within the temperate zone. However, past stability in migratory populations does not extend to the present day. Ecological monitoring data clearly show that contemporary populations of long-distance migrants are facing steep decline (54). Our discovery of association between genetic diversity and migration distance provides important context for understanding the situation faced by migratory populations during rapid environmental change. The relatively high genetic diversity displayed by long-distance migratory species could potentially help their populations adapt to new conditions (33,55), but many of these populations will also have to persist through population declines on a scale that may have little precedent in their recent evolutionary histories.

## Materials and Methods

### Species and sampling

Our study system includes 35 co-distributed boreal forest bird species (table S1, Fig. S36). These species represent a core subset of the small-bodied species (< 75 grams) that breed widely across the boreal forest and the temperate-boreal transition (hemiboreal) habitat of our sampling area (corresponding to the “Northern Forests” Level I Ecoregion; 18). Although all 35 species we sampled are found sympatrically during the breeding season throughout much of the boreal ecoregion, they exhibit variation in microhabitat preference and the extent of their geographic range beyond the boreal forest ecoregion (12,18,56,57). Three species are woodpeckers (Piciformes), and the remaining are from 17 genera and 10 families of songbirds (Passeriformes). We focus on obligate migratory species and nonmigratory species, meaning we did not include species primarily characterized by nomadic or irruptive movements (e.g., Fringillidae, Sittidae, and Bombycillidae) in our analyses. Our study species vary greatly in migratory strategy and distance but share many life history attributes (e.g., mating system and age at first breeding season; 19).

We sampled species broadly and evenly along the longitudinal axis of this ecoregion (Fig. 1), from Alberta, Canada east of the Rocky Mountains in the west to the mainland Maritime Provinces of Canada (Quebec, New Brunswick, and Nova Scotia) in the east, as well as intervening locations in central Canada (Manitoba) and the northern United States (northern Michigan, Minnesota, and New York). Our sampling was intentionally designed to capture continuous spatial variation across the eastern boreal region shared broadly by the species in our study. We did not sample populations in regions at the extreme edges of the boreal region (Alaska, Newfoundland) or in other ecoregions (e.g. the Rocky Mountains) because not all species’ ranges extend into these regions, and many species distributed across these ecoregions have experienced phylogeographic breaks between them (13–15,58–64). All samples were collected during the breeding season. We sequenced DNA from 1673 samples (table S1; mean = 47 ± 13 samples per species, range = 15–67 samples). Most (88.3%) of the samples were ethanol-preserved or flash-frozen specimen-vouchered tissues provided by contributing natural history collections (data S1), while the remaining 11.7% came from unvouchered blood samples. Field sampling was approved by the University of Michigan Institutional Animal Care and Use Committee and all relevant permitting authorities (see Acknowledgements) and performed in accordance with the Ornithological Council’s Guidelines for Use of Wild Birds in Research (65).

### Preparation of DNA sequence data

We extracted DNA using DNeasy Blood and Tissue Kits (Qiagen Sciences, Germantown, MD, USA) or phenol-chloroform extraction. Libraries were prepared using a modified Illumina Nextera library preparation protocol (66,67) and then sequenced on either an Illumina HiSeq or Illumina NovaSeq 6000 platform using paired-end sequencing of 150 bp reads. Libraries were prepared using a modified Illumina Nextera library preparation protocol (66,67) and then sequenced on an Illumina HiSeq, NovaSeq 6000, or NovaSeq X platform using paired-end sequencing of 150 bp reads. Libraries used in analyses produced an average of 32 million ± 10 million reads per sample (range 7 million–156 million).

We trimmed remaining adaptors and low-quality bases from demultiplexed data with AdapterRemoval v2.3.1 using the –trimns and –trimqualities options (68). To mitigate potential batch effects associated with differences between the NovaSeq and HiSeq platforms, we used fastp v0.23.2 (69) with the--cut_right option to remove low-quality read ends (70). Following this filter, the mean number of bases per individual at a base quality of at least 30 was 4.3 billion ± 1.3 billion bases (range: 0.94 billion–21.2 billion).

We aligned samples to a reference genome from a related species (71–78) (data S2) using bwa mem (79) and then sorted them using SAMtools (80). We removed overlapping reads using clipOverlap in bamUtil (81), then marked duplicate reads with MarkDuplicates and assigned all reads to a new read-group with AddOrReplaceReadGroups using picard (http://broadinstitute.github.io/picard/). All bam alignment files were then indexed using SAMtools. Samples had an average duplication rate of 7% (sd 4%, range 0.3% to 33%).

Excluding duplicate reads, mean mapping rate was 88% (sd 7%, range 38% to 98%). Finally, we re-aligned samples around indels using GATK v3.7 (82). We applied the GATK RealignerTargetCreator tool to the entire dataset analyzed for each species and applied the GATK IndelRealigner tool to each sample. We calculated the average genome-wide coverage for each alignment by dividing the total coverage of the bam file (calculated with the SAMtools depth function, filtered to exclude reads mapped with a quality lower than 30) by the total length of the reference genome. Bases with a coverage of 0 (including segments of the reference genome where no reads were mapped) are included in these calculations, making them conservative. Across all aligned samples, average genome-wide coverage of high-quality mapped reads was 2.8x (± 1x).

### Data quality control using mitochondrial sequences

We confirmed species identity by examining at least one mitochondrial gene per successful assembly via BLAST (https://blast.ncbi.nlm.nih.gov/Blast.cgi) in Geneious (v. 2021.2.2). Mitochondrial genomes were assembled using NOVOplasty v4.3.1 (83) and analysed as described in (38). We also visually inspected alignments of mitochondrial genes to check for evidence of cross-contamination and hybridization. Based on these assessments, we excluded 14 samples with an unexpected mitochondrial species identity due to specimen misidentification or hybridization, as well as 5 samples with evidence of mitochondrial chimerism, which is likely due to sample cross-contamination. None of these excluded samples are counted in sample totals reported elsewhere or included in downstream analyses.

### Data filtering

We used ANGSD v0.941(84) to calculate genotype likelihoods for all sites inferred by ANGSD to be SNPs at an alpha-level 0.05. We applied ngsParalog v1.3.2 (85) to filter out SNPs with a high likelihood of occurring within a mis-mapped or paralogous region. Next, we used PCAngsd v1.10 (86) to create a PCA separately for each chromosome. We observed many chromosomes with evidence of PCA clustering consistent with the presence of inversion polymorphisms (87,88), so we analyzed each chromosome further using lostruct (89) as implemented using PCAngsd with scripts available from https://github.com/alxsimon/local_pcangsd. Inversion polymorphisms as well as polymorphisms in large linked haplotypes associated with other forms of suppressed recombination can obscure spatial genetic patterns (40,90), so we removed all such regions from each species’ dataset (data S3). We flagged regions as polymorphisms when they met two criteria: 1) Lostruct identified the region as showing distinct local structure, and 2) PCA with data from the region showed distinct clustering into two or three groups, indicating a significant polymorphism within the species.

None of the regions we flagged with these criteria showed strong spatial structure. We removed entire microchromosomes (<35 Mb in length; 91) with flagged regions on them. To remove flagged regions from macrochromosomes without discarding the entire chromosome, we used lostruct plots to identify the affected regions and we removed the region with a buffer of at least 1 Mb on either side. When macrochromosomes showed evidence of more than one flagged region and/or if the affected region was not clearly identifiable using lostruct, we discarded the entire chromosome (data S3). Finally, we removed all reads mapped to sex chromosomes or unassembled scaffolds from our dataset.

We also used chromosome-level PCA plots to further filter individuals. We removed individuals if they presented as a strong PCA outlier on multiple macrochromosomes. In all three of the sparrow species and four warbler species (*Leiothlypis peregrina, Leiothlypis ruficapilla, Setophaga castanea, Setophaga magnolia*), we detected sex-based clustering on autosomes as well as sex chromosomes. We filtered females out of these species because we have many more samples from males than females. Individuals filtered out at this stage are not included in the sample numbers we report here.

### Estimation of spatial genetic structure within each species

After examining the data and applying the quality filters described above, we then used PCAngsd v1.10 (86) with the –admix option to create full-genome PCAs and admixture plots for each species. We plotted the results of these analyses in R. PCAngsd also produces a matrix of covariances between each pair of individuals for each species, which we used in downstream modeling of isolation by distance. We estimated isolation by distance using individual-based genetic covariance instead of more-typical *F_st_* estimates because the latter require *a priori* definition of populations, which is not feasible within our continuous sampling area. For each pair of samples, we calculated geographic distance using the function ‘distGeo’ from the R package ‘geosphere’ (92) on latitude/longitude coordinates.

### Estimation of species attributes hypothesized to affect spatial structure

We created a species-level estimate for each of the attributes we hypothesized to influence spatial structure (seasonal migration, mass, habitat association, and genetic diversity; data S1). We represented variation in migratory behavior both as a binary variable (migratory vs nonmigratory) and as a continuous estimate of migration distance, as represented by the distance between the centroids of species’ breeding and wintering ranges (19,93). Mean mass estimates for each species are from (94,95). We used a binary variable to indicate whether species show association with early successional habitat based on our field experience with these species and information from (95).

### Estimation of genetic diversity using subsampled loci

To estimate genetic diversity (θ_π_), we created a subsampled set of genome loci for each species using scripts modified from https://github.com/markravinet/genome_sampler. Each subset was created by sampling random 2kb loci at least 10kb apart, which produces loci comprising roughly 10% of the genome (data S2). Regions removed by the inversion filters described above were excluded prior to random locus selection. After locus selection, we removed SNPs from the selected loci that were flagged by ngsParalog as described above. Individuals removed from PCA-based filters described above were excluded. The subsampled loci were stored in a BED file and supplied as a site filter to ANGSD in the next step. To estimate θ_π_, we created a species-wide SAF file in ANGSD and used the program winsfs v0.7.0 (96) to generate and fold a species-wide 1-d SFS. We then used ANGSD’s saf2theta and thetaStat functions on the SAF and SFS to generate estimates of θ_π_ (specifically, θ_π_ of 97). To calculate genome-wide θ_π_, we divided the sum of θ_π_ across all loci by the sum of the number of sites analyzed per locus.

### Hierarchical modeling of the effects of traits on spatial structure

We used the R package ‘HMSC’ (39) to jointly estimate spatial structure for each species and the effects of species’ traits, or attributes, on these patterns at the community level. This approach has advantages over analogous two-step approaches (those that model slopes generated from one model as the outcome variable in a second model) because hierarchical modeling propagates error across the levels of the model, making it ideal for use in this context (e.g., 98). We model spatial structure as a linear relationship between pairwise genetic covariance (calculated in PCAngsd) and pairwise geographic distance (of sampling location) for all conspecific pairs of samples in our dataset. These relationships are represented as *β* coefficients (which we refer to as *β_IBD_*) for each species in the HMSC model. The effect of each attribute on interspecific variation in *β_IBD_* is represented by the attribute’s γ coefficient in the HMSC model. We provided the HMSC model with a phylogenetic tree of our study species prepared using data from birdtree.org (99), following (100) as described in (35,93) (Fig. S36). We centered and scaled all traits prior to modeling to have a mean of 0 and a standard deviation of 1.

To test whether migratory status (migratory vs nonmigratory) affects spatial structure differently than variation in migration distance, we created two HMSC models. The first includes all 35 study species and includes migratory status as a binary trait in addition to the five other traits we investigated. The second includes only the 30 migratory species in our study and represents migration behavior using our continuous estimate of migration distance.

We ran HMSC models with the options ‘YScale = T’ and ‘distr = “normal”’ and all other options at the default. We used the HMSC’s default priors, which are designed to be generally applicable and are described in detail in (46). We ran the models in four replicate Markov Chain Monte Carlo (MCMC) chains. After a 10,000 iteration burn-in, we let the chains run for 75,000 iterations and thinned the output by 50, producing 1500 samples per chain from the posterior distribution for each parameter (98). We examined trace plots and effective sample sizes to ensure model convergence and adequate independence of samples.

We examined and visualized the posterior estimates and posterior support values for each model parameter of interest (using functions from the R packages HMSC, MCMCvis (101), and bayesplot (102), which create summaries of model output across all four chains. We estimated trait R^2^ values with the HMSC function ‘computeVariancePartitioning().’ We performed variance partitioning on 250 posterior samples from our models, as opposed to all 1500 used in other analyses, for computational feasibility.

### Assessing the relationship between genetic diversity and migration distance

We used the function ‘pgls()’ from the R package ‘caper’ (103) to assess the relationship between genetic diversity and migration distance in a phylogenetic framework. We scaled the variables to have a mean of 0 and a standard deviation of 1 prior to modeling and we supplied the function with the same phylogenetic tree we used in the HMSC models, described above. We used the option ‘lambda=“ML”,’ which causes the model to estimate a maximum likelihood estimate of λ (phylogenetic signal in model residuals).

## Acknowledgments

For comments and advice, we thank G. Bradburd, R. Burner, K. Wacker, E. Gulson-Castillo, V. Gomez-Bahamon, J. Berv, Z. Hancock, M. Hack, A. Marshall, L. Knowles, D. Rabosky, M. Witynksi, and A Benavides. J. Megahan provided artwork for Fig. 1. For providing samples, we thank the curators, collections staff, and field collectors from the following institutions: American Museum of Natural History, Cleveland Museum of Natural History, Cornell University Museum of Vertebrates, New York State Museum, Royal Alberta Museum, Royal Ontario Museum, University of Alaska Museum of the North, University of Minnesota Museum of Natural History, University of California, Berkeley Museum of Vertebrate Zoology, University of Michigan Museum of Zoology, University of Washington Burke Museum. For additional samples, we thank J. Tremblay (Environment and Climate Change Canada). For field permits, we thank the United States Fish and Wildlife Service, the United States Forest Service, the Minnesota Department of Natural Resources, the Michigan Department of Natural Resources, the Canadian Wildlife Service of Environment and Climate Change Canada, Alberta Fish and Wildlife and Manitoba Fish and Wildlife. Field sampling was approved by the University of Michigan Animal Care and Use Committee (# PRO00010608). For assistance in the field, we thank C. Brennan, S. Campbell, S. DuBay, G. M. Erickson, M. M. Ferraro, A. M. FitzGerald, L. Gooch, E. Gulson-Castillo, J. Ralston, C. Scobey, H. Skeen, V. Ting, K. Wacker, and members of the Burg lab. For assistance in the lab, we thank T. Schweizer, C. Rayne, R. Herman, J. Yan, C. Kaczmarek, M. Florkowski, M. Guza, C. Pajka, M. Hack, and C. Jordan. Next-generation sequencing for this project was partially carried out in the Advanced Genomics Core at the University of Michigan. This research was also supported in part through computational resources and services provided by Advanced Research Computing (ARC), a division of Information and Technology Services (ITS) at the University of Michigan, Ann Arbor.

## Funding

National Science Foundation grant 2146950 (BMW)

National Science Foundation Graduate Research Fellowship DGE 1256260, Fellow ID 2018240490 (TMP)

Jean Wright Cohn Endowment Fund at the University of Michigan Museum of Zoology

Robert W. Storer Endowment Fund at the University of Michigan Museum of Zoology

Mary Rhoda Swales Museum of Zoology Research Fund at the University of Michigan Museum of Zoology

William G. Fargo Fund at the University of Michigan Museum of Zoology

William A. and Nancy R. Klamm Endowment funds at the Cleveland Museum of Natural History

University of Michigan Rackham Graduate Student Research Grant

## Notes

### Competing Interest Statement

The authors have declared no competing interest.

