## Supplemental Tables, Figures, Captions for "Population genetic consequences of the seasonal migrations of birds"

Supplementary materials for **Population genetic consequences of the seasonal migrations of birds**

**Fig. S1 to S35. (separate file)**

Spatial genetic patterns for each of the 35 species in the study are shown with PCA **(A)**, and admixture **(B)** plots, where point colors correspond to panel **C**, which shows the distribution of sampling for each species. Panel **D** shows *β_IBD_* plots with a line showing the slope and intercept estimated by the HMSC model with all species.

**Fig. S36.**

The phylogenetic relationship among the species in the study is shown on the tree from birdtree.org used in our analyses. The tree is plotted alongside a heatmap representing interspecific variation in the four species attributes we investigated in HMSC models (migration distance, mass, association with early successional habitat, and genetic diversity).

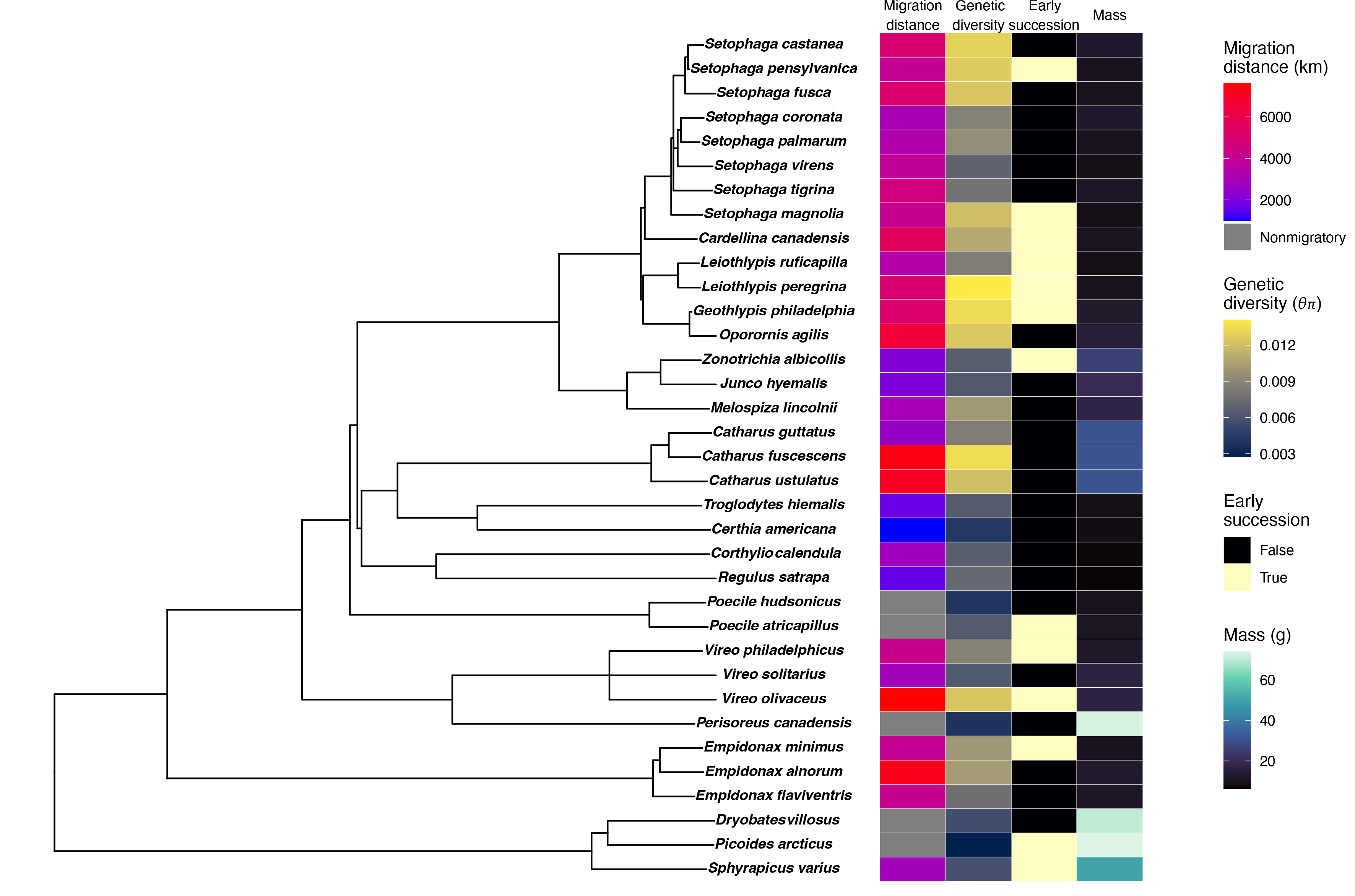

**Fig. S37.**

The relationship between migration distance and time limitation on the breeding grounds among migratory species (*n*=30), showing that long-distance migrants have shorter breeding seasons in the boreal region. This figure is adapted from Supplementary Figure 2 of (*5*).

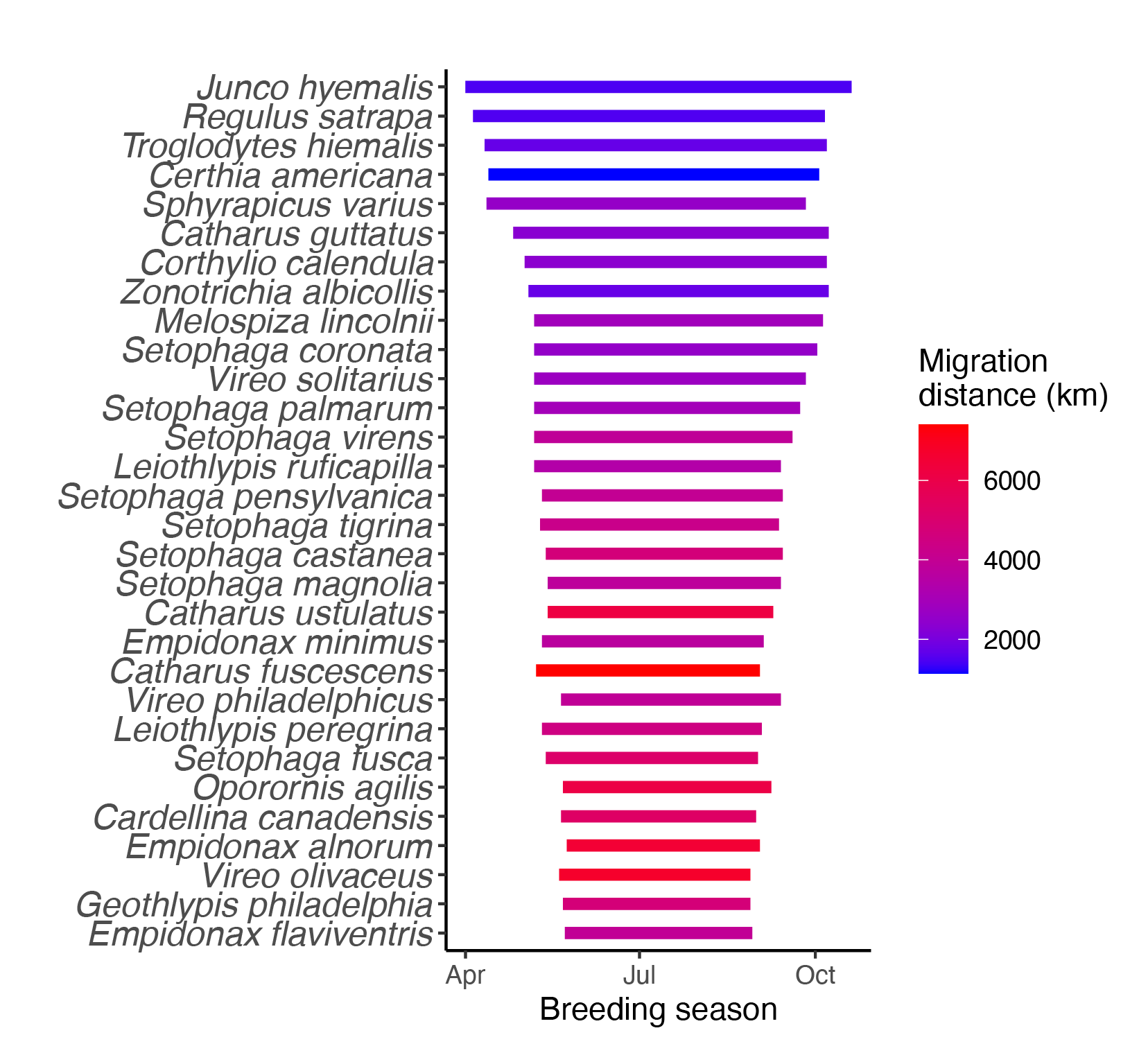

| Species | Family | Sample number | Migration distance (km) | Migratory status | Mean mass (g) | Early successional habitat association | Genetic diversity (*θ_π_*) | *β_IBD_* | β_IBD_ posterior support |
| --- | --- | --- | --- | --- | --- | --- | --- | --- | --- |
| *Picoides arcticus* | Picidae | 31 | 0 | FALSE | 74 | TRUE | 0.0027 | -0.1083 | 0.9985 |
| *Dryobates villosus* | Picidae | 43 | 0 | FALSE | 70.36 | FALSE | 0.0056 | -0.6978 | 1.0000 |
| *Sphyrapicus varius* | Picidae | 62 | 2900 | TRUE | 50.3 | TRUE | 0.0058 | -0.4782 | 1.0000 |
| *Empidonax alnorum* | Tyrannidae | 49 | 7300 | TRUE | 12.79 | FALSE | 0.0102 | -0.0223 | 0.8723 |
| *Empidonax flaviventris* | Tyrannidae | 58 | 4100 | TRUE | 11.6 | FALSE | 0.0077 | -0.1635 | 1.0000 |
| *Empidonax minimus* | Tyrannidae | 44 | 4000 | TRUE | 10.3 | TRUE | 0.0099 | -0.0291 | 0.8387 |
| *Perisoreus canadensis* | Corvidae | 25 | 0 | FALSE | 73 | FALSE | 0.0039 | -0.6740 | 1.0000 |
| *Vireo olivaceus* | Vireonidae | 63 | 7600 | TRUE | 16.7 | TRUE | 0.0124 | -0.2431 | 1.0000 |
| *Vireo philadelphicus* | Vireonidae | 17 | 4200 | TRUE | 12.2 | TRUE | 0.0088 | 0.0248 | 0.3338 |
| *Vireo solitarius* | Vireonidae | 56 | 2900 | TRUE | 16.6 | FALSE | 0.0064 | -0.4373 | 1.0000 |
| *Poecile atricapillus* | Paridae | 59 | 0 | FALSE | 10.8 | TRUE | 0.0064 | -0.7316 | 1.0000 |
| *Poecile hudsonicus* | Paridae | 37 | 0 | FALSE | 9.8 | FALSE | 0.0041 | -0.7770 | 1.0000 |
| *Corthylio calendula* | Regulidae | 49 | 2800 | TRUE | 6.68 | FALSE | 0.0067 | -0.0597 | 0.9960 |
| *Regulus satrapa* | Regulidae | 63 | 1600 | TRUE | 6.23 | FALSE | 0.0073 | -0.4849 | 1.0000 |
| *Certhia americana* | Certhiidae | 48 | 1000 | TRUE | 8.4 | FALSE | 0.0043 | 0.0154 | 0.2773 |
| *Troglodytes hiemalis* | Troglodytidae | 26 | 1700 | TRUE | 8.9 | FALSE | 0.0064 | 0.0116 | 0.4107 |
| *Catharus fuscescens* | Turdidae | 47 | 7400 | TRUE | 31.2 | FALSE | 0.0134 | -0.8952 | 1.0000 |
| *Catharus guttatus* | Turdidae | 65 | 2500 | TRUE | 31 | FALSE | 0.0085 | -0.0660 | 1.0000 |
| *Catharus ustulatus* | Turdidae | 66 | 7100 | TRUE | 30.8 | FALSE | 0.0121 | -0.3142 | 1.0000 |
| *Junco hyemalis* | Passerellidae | 59 | 2000 | TRUE | 19.89 | FALSE | 0.0063 | -0.0429 | 0.9947 |
| *Melospiza lincolnii* | Passerellidae | 53 | 3000 | TRUE | 17.4 | FALSE | 0.0101 | -0.0581 | 0.9988 |
| *Zonotrichia albicollis* | Passerellidae | 68 | 2100 | TRUE | 25.9 | TRUE | 0.0065 | -0.0303 | 0.9768 |
| *Cardellina canadensis* | Parulidae | 31 | 5500 | TRUE | 10.42 | TRUE | 0.0109 | -0.6831 | 1.0000 |
| *Geothlypis philadelphia* | Parulidae | 52 | 5100 | TRUE | 12.61 | TRUE | 0.0134 | -0.8201 | 1.0000 |
| *Oporornis agilis* | Parulidae | 30 | 6600 | TRUE | 15.2 | FALSE | 0.0125 | 0.0442 | 0.0995 |
| *Leiothlypis peregrina* | Parulidae | 47 | 5000 | TRUE | 10.02 | TRUE | 0.0141 | 0.0188 | 0.1660 |
| *Leiothlypis ruficapilla* | Parulidae | 57 | 3400 | TRUE | 8.73 | TRUE | 0.0085 | -0.0937 | 1.0000 |
| *Setophaga castanea* | Parulidae | 45 | 5000 | TRUE | 12.59 | FALSE | 0.0130 | -0.0172 | 0.8332 |
| *Setophaga coronata* | Parulidae | 70 | 3100 | TRUE | 12.51 | FALSE | 0.0088 | -0.0880 | 1.0000 |
| *Setophaga fusca* | Parulidae | 52 | 5100 | TRUE | 9.7 | FALSE | 0.0124 | -0.1667 | 1.0000 |
| *Setophaga magnolia* | Parulidae | 56 | 4100 | TRUE | 8.72 | TRUE | 0.0120 | -0.4074 | 1.0000 |
| *Setophaga palmarum* | Parulidae | 51 | 3300 | TRUE | 10.3 | FALSE | 0.0094 | -0.5598 | 1.0000 |
| *Setophaga pensylvanica* | Parulidae | 46 | 4000 | TRUE | 9.64 | TRUE | 0.0127 | -0.0813 | 0.9997 |
| *Setophaga tigrina* | Parulidae | 44 | 4700 | TRUE | 11 | FALSE | 0.0079 | 0.0017 | 0.4892 |
| *Setophaga virens* | Parulidae | 63 | 3900 | TRUE | 8.8 | FALSE | 0.0069 | -0.5739 | 1.0000 |

**Table S1.**

Species used in analyses along with their taxonomic family and the number of samples used, values for species attributes used in HMSC models to infer γ coefficients, and posterior estimates of *β_IBD_* and its mean posterior probability from the version of our HMSC model that included all species. *β_IBD_* estimates from the other version of the HMSC model (without nonmigratory species) were highly similar.

**Data S1. (separate file)**

List of samples used and metadata for each sample, including species and sex, sample numbers, providing institutions, tissue types, and the date and locality where the sample was collected. We also show bioinformatic metadata for each sample (see also Methods), including the total number of 150-bp paired-end reads per sample produced during sequencing, duplication rate, the total number of base pairs sequenced (in billions), the number of base pairs with a quality of 30 or higher (in billions), and percent of reads (after filtering) that mapped to each sample’s respective reference genome (Supplementary Table 2). The columns indicating numbers of base pairs sequenced provide an approximation of coverage depth for each sample. Museum abbreviations: AMNH=American Museum of Natural History; CMNH=Cleveland Museum of Natural History; CUMV=Cornell University Museum of Vertebrates; MMNH=Bell Museum of Natural History, University of Minnesota; MVZ=Museum of Vertebrate Zoology, University of California Berkeley; NYSM=New York State Museum; RAM=Royal Alberta Museum; ROM=Royal Ontario Museum; UAM=University of Alaska Museum of the North; UMMZ=University of Michigan Museum of Zoology; UWBM=University of Washington Burke Museum

**Data S2. (separate file)**

Bioinformatic metadata for each species, including GenBank accession numbers of reference genomes and the numbers of loci used in bioinformatic analyses. These numbers reflect how many loci were analyzed after filtering. Variation in the outcome of filtering (e.g. the number of regions excluded based on filtering of inversion polymorphisms) is responsible for some interspecific variation in these numbers, particularly in the length of the genome subset used to estimate θπ. The number of SNPs for each species also varies according to the species’ genetic diversity.

**Data S3. (separate file)**

Information about chromosomes and large genomic regions excluded from analyses based on evidence for possible inversion polymorphism. Each filtered region is listed along with the species for which it was filtered, the GenBank accession name of the affected chromosome, and a column indicating whether the entire chromosome was excluded from analyses. In cases where only part of the chromosome was excluded, we list the start and end of the excluded region.
