## Supplementary material for "Population genetic consequences of the seasonal migrations of birds": Figs S1 to S35

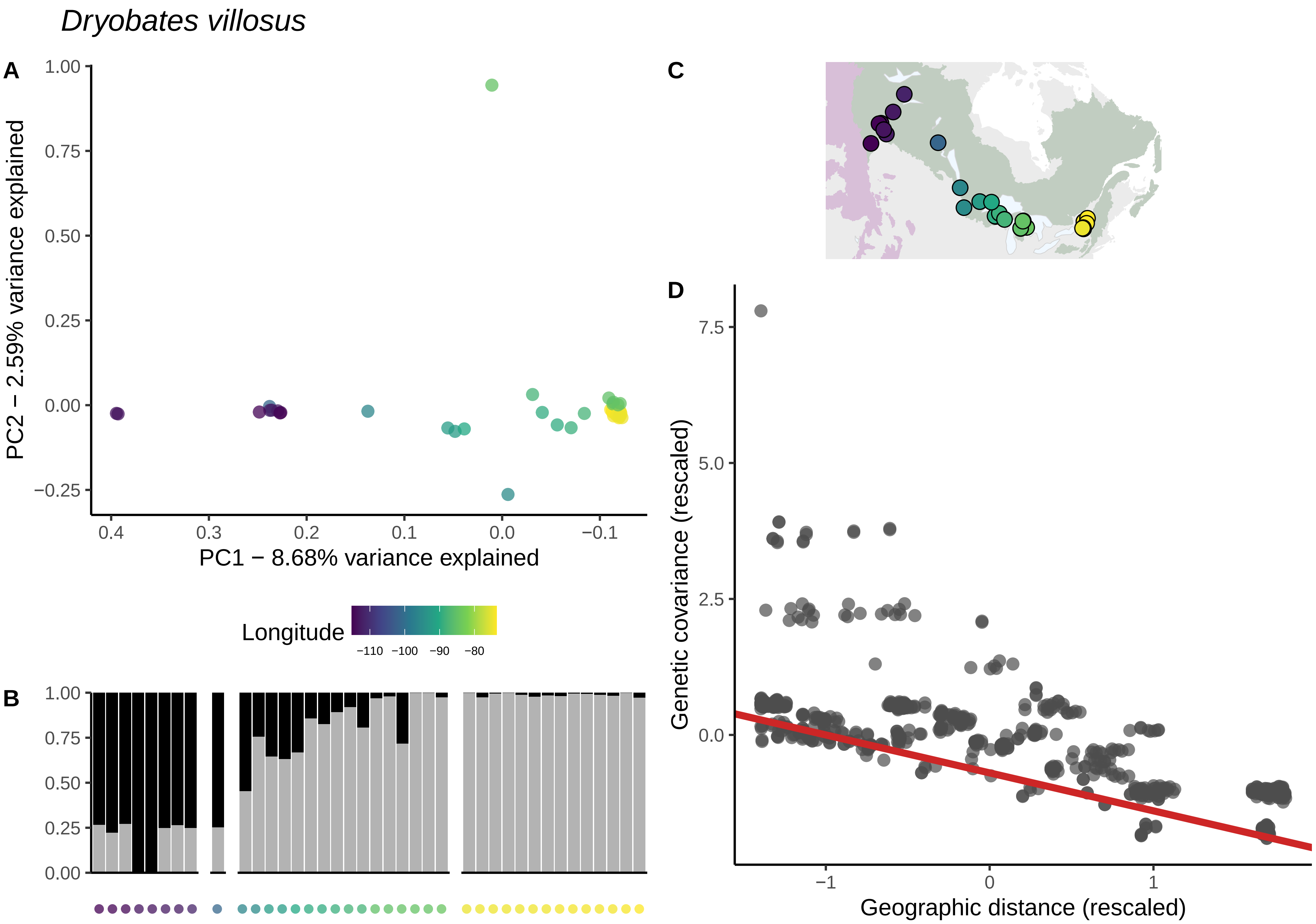
Dryobates villosus
A
PC2 − 2.59% variance explained
PC1 − 8.68% variance explained
Longitude
−110 −100 −90 −80
B
C
D
Genetic covariance (rescaled)
Geographic distance (rescaled)

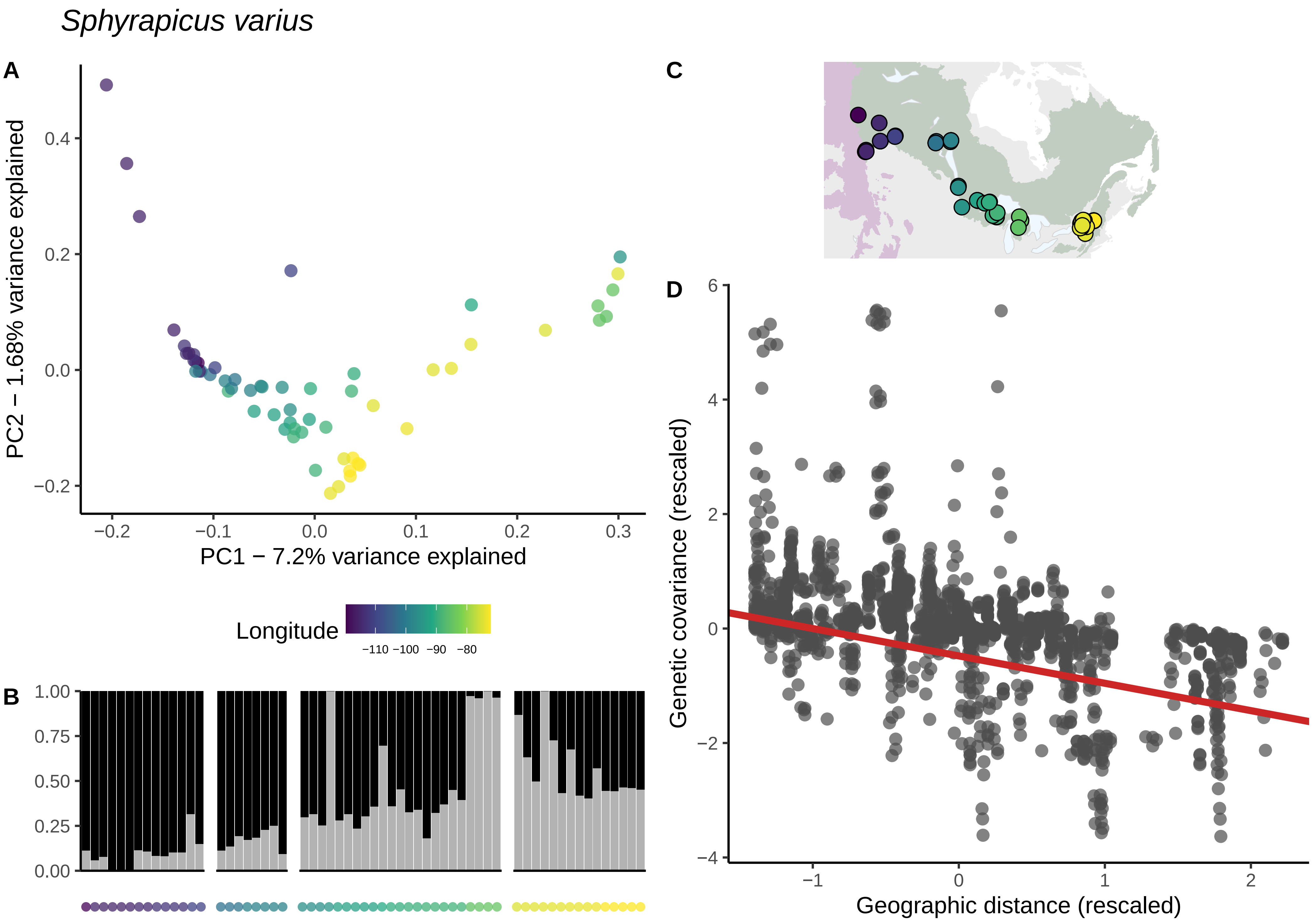

Sphyrapicus varius
A
PC2 − 1.68% variance explained
PC1 − 7.2% variance explained
0.4
0.2
0.0
−0.2
−0.2
−0.1
0.0
0.1
0.2
0.3
Longitude
−110 −100 −90 −80
B
1.00
0.75
0.50
0.25
0.00
C
D
Genetic covariance (rescaled)
Geographic distance (rescaled)
6
4
2
0
−2
−4
−1
0
1
2

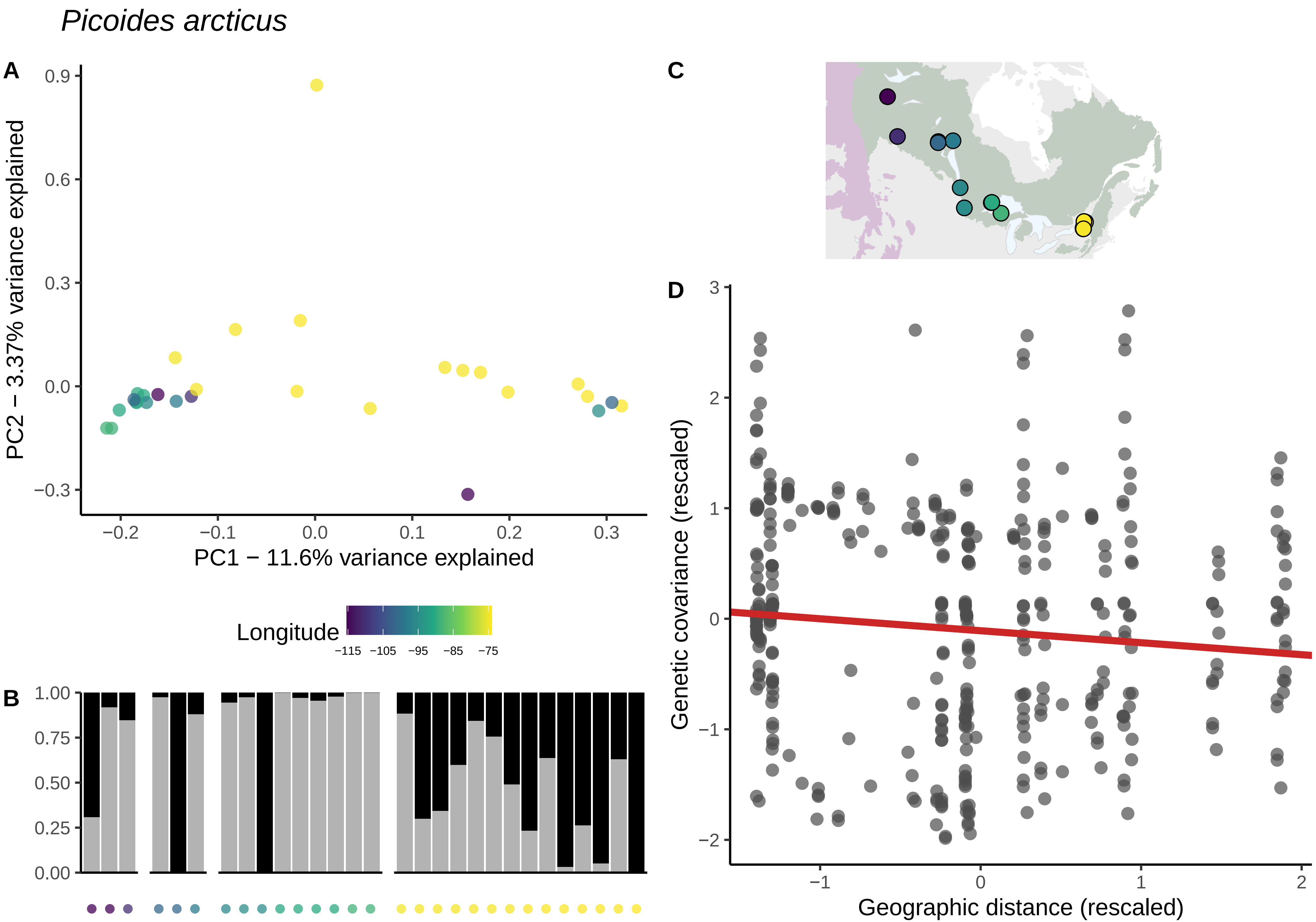

Picoides arcticus
A
PC2 − 3.37% variance explained
PC1 − 11.6% variance explained
Longitude
−115 −105 −95 −85 −75
B
C
D
Genetic covariance (rescaled)
Geographic distance (rescaled)

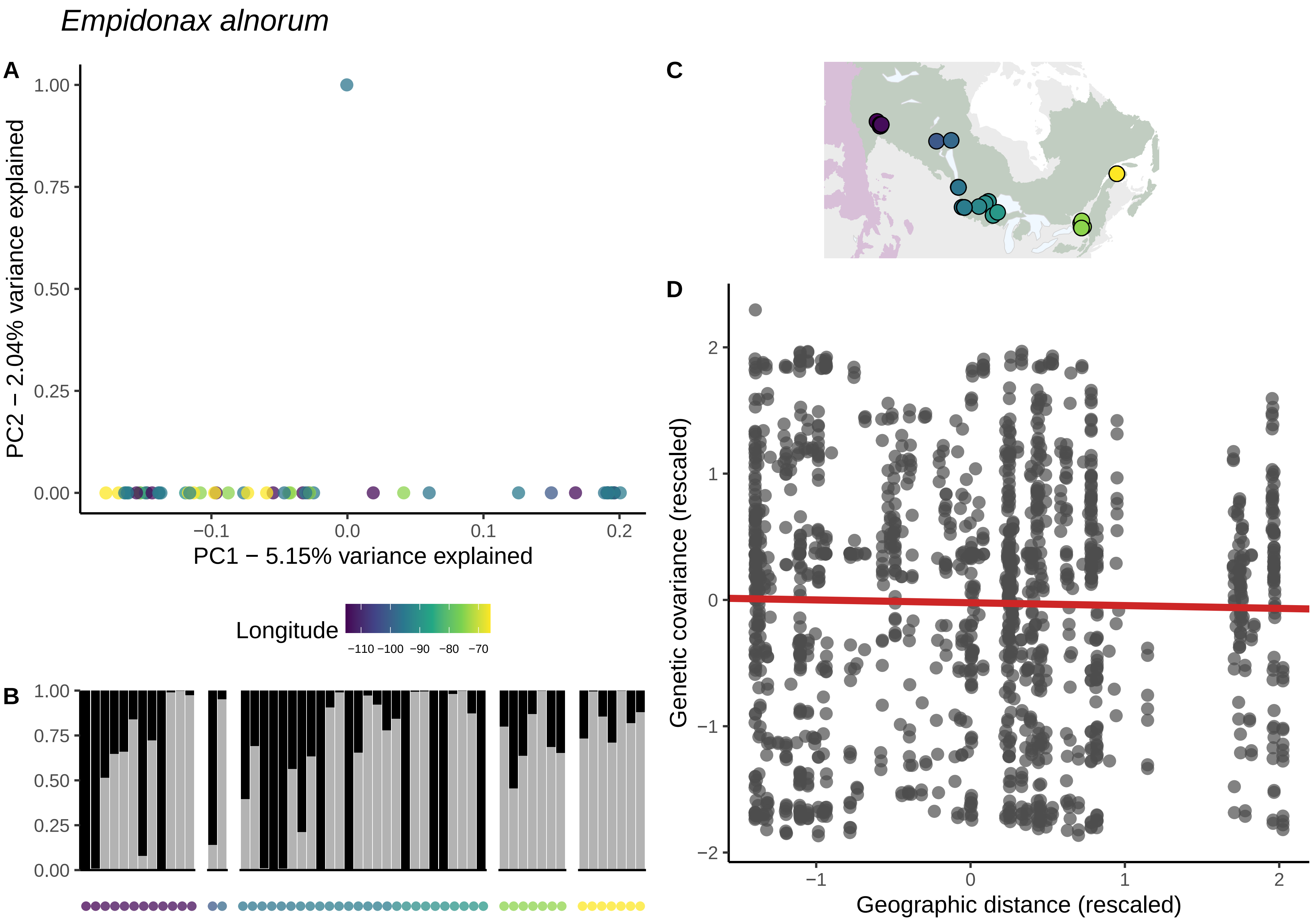

Empidonax alnorum
A
PC2 − 2.04% variance explained
PC1 − 5.15% variance explained
1.00
0.75
0.50
0.25
0.00
−0.1
0.0
0.1
0.2
Longitude
−110 −100 −90 −80 −70
B
1.00
0.75
0.50
0.25
0.00
C
D
Genetic covariance (rescaled)
Geographic distance (rescaled)
2
1
0
−1
−2

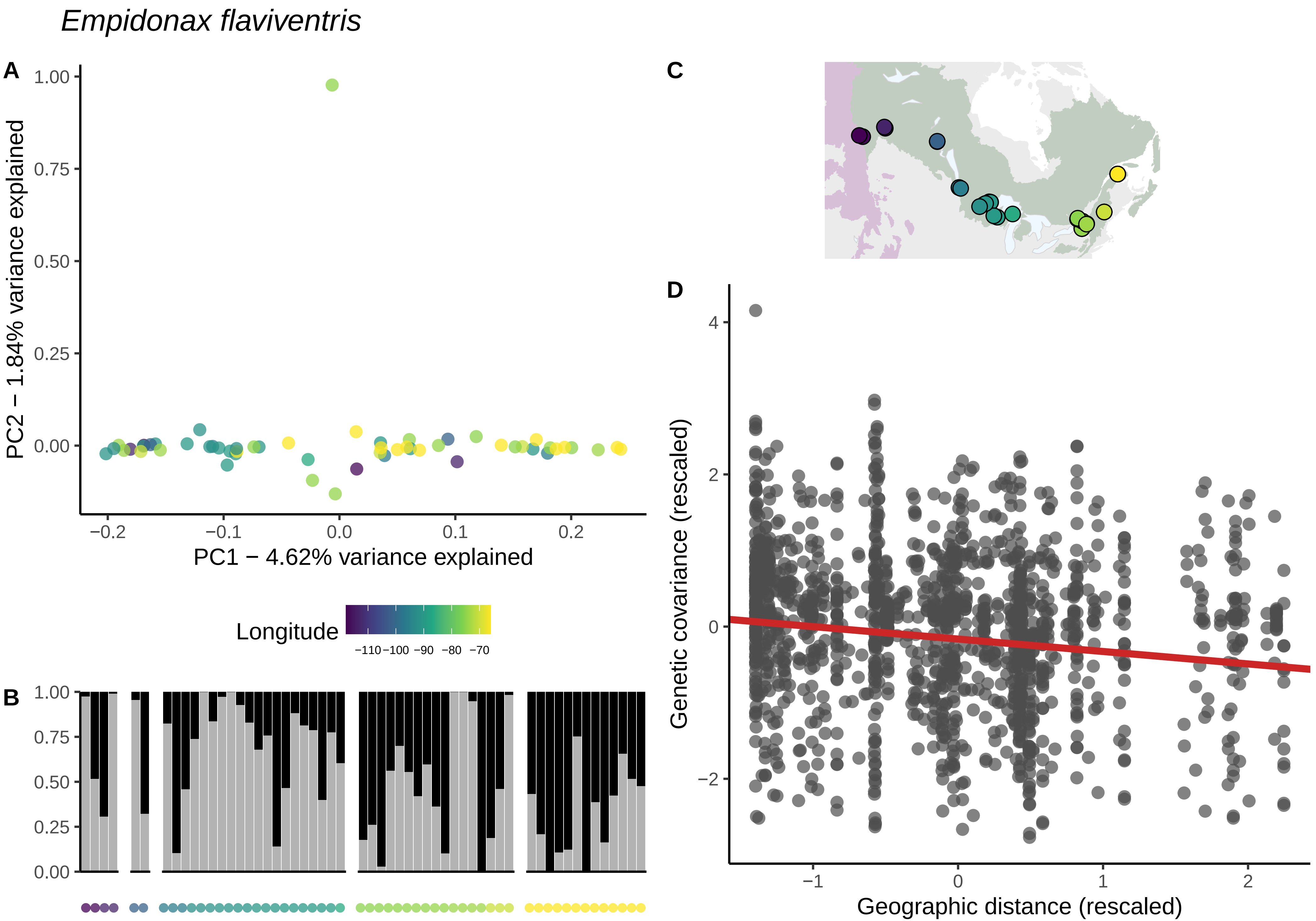

Empidonax flaviventris
A
B
C
D
PC2 − 1.84% variance explained
PC1 − 4.62% variance explained
1.00
0.75
0.50
0.25
0.00
−0.2
−0.1
0.0
0.1
0.2
Longitude
−110
−100
−90
−80
−70
Genetic covariance (rescaled)
Geographic distance (rescaled)
4
2
0
−2
−1
1

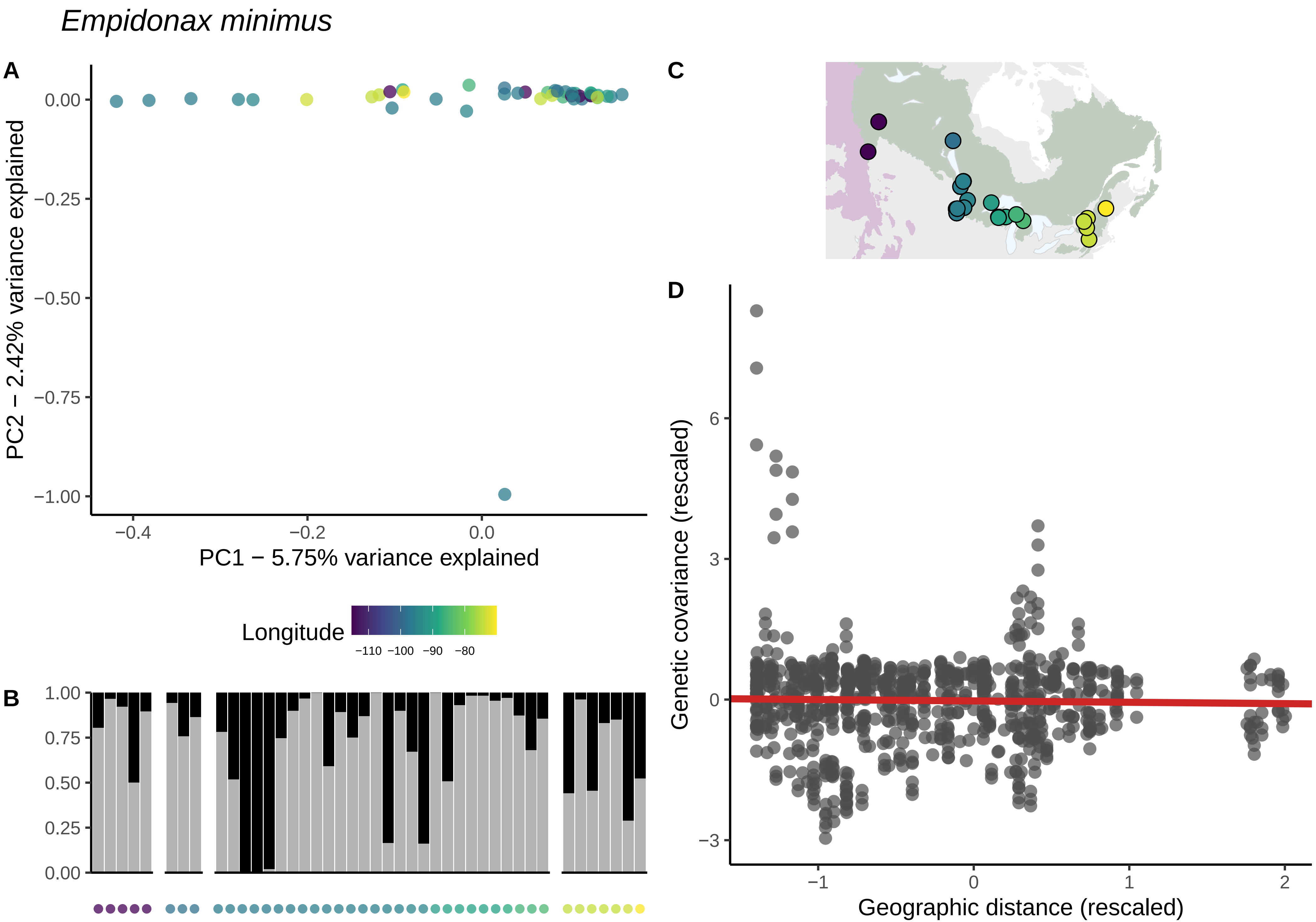

*Empidonax minimus*
A
B
C
D
PC2 − 2.42% variance explained
PC1 − 5.75% variance explained
0.00
−0.25
−0.50
−0.75
−1.00
−0.4
−0.2
0.0
Longitude
−110 −100 −90 −80
1.00
0.75
0.50
0.25
0.00
Genetic covariance (rescaled)
Geographic distance (rescaled)
6
3
0
−3
−1
0
1
2

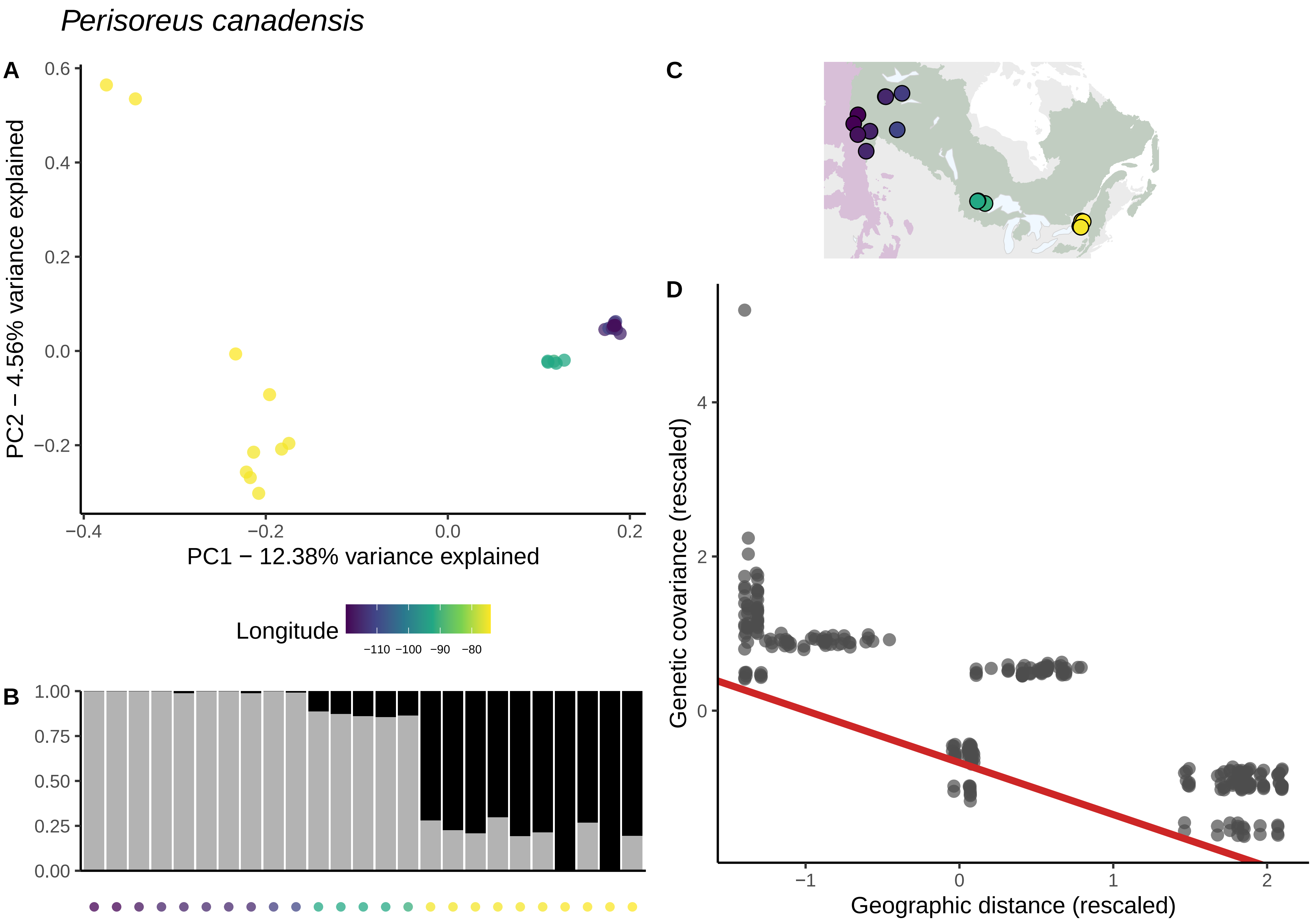
Perisoreus canadensis
A
PC2 − 4.56% variance explained
PC1 − 12.38% variance explained
0.6
0.4
0.2
0.0
−0.2
−0.4
−0.2
0.0
0.2
Longitude
−110 −100 −90 −80
B
1.00
0.75
0.50
0.25
0.00
C
D
Genetic covariance (rescaled)
Geographic distance (rescaled)
4
2
0
−1
0
1
2

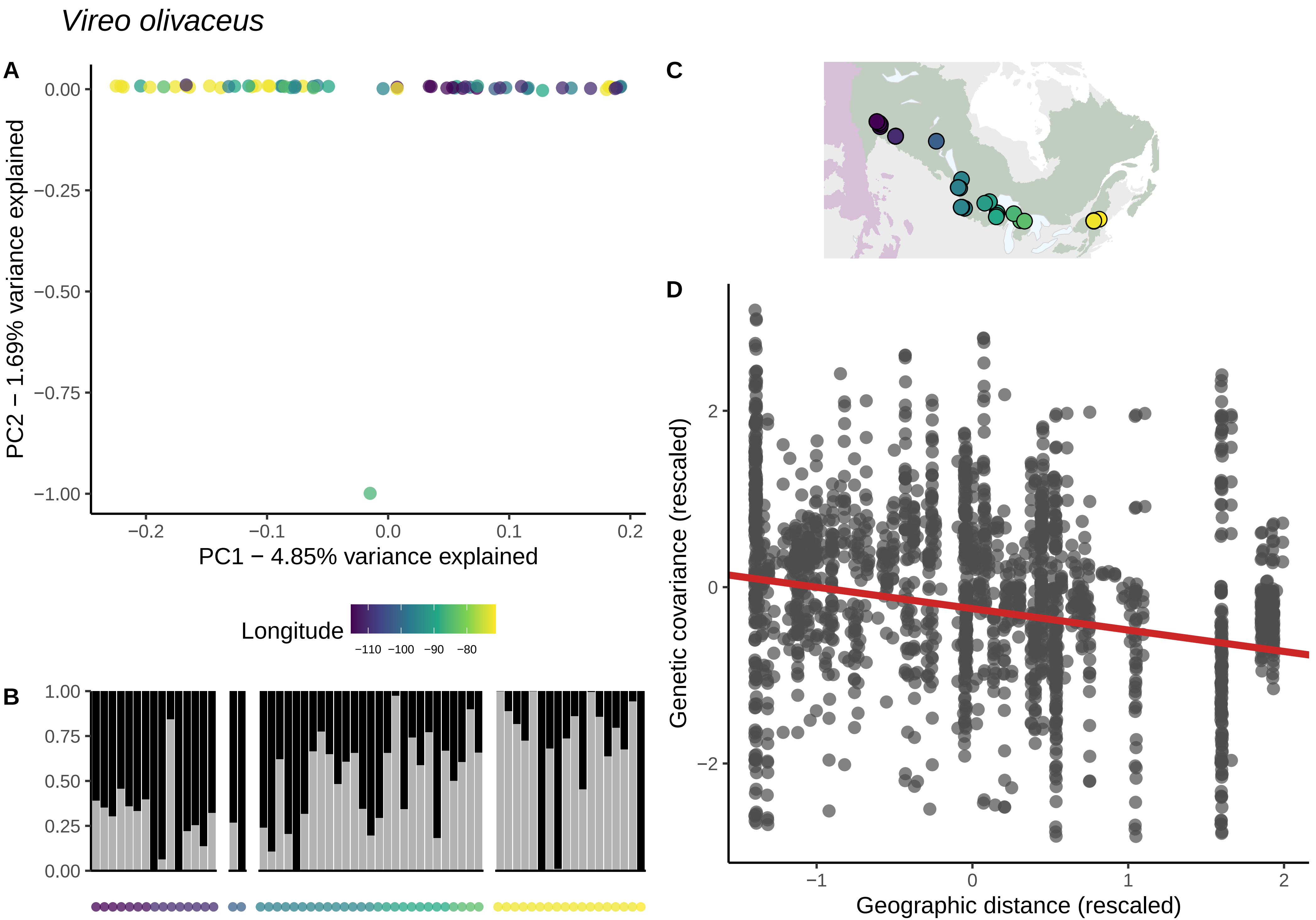
Vireo olivaceus
A
0.00
−0.25
−0.50
−0.75
−1.00
PC2 − 1.69% variance explained
−0.2
−0.1
0.0
0.1
0.2
PC1 − 4.85% variance explained
Longitude
−110 −100 −90 −80
B
1.00
0.75
0.50
0.25
0.00
C
D
2
0
−2
Genetic covariance (rescaled)
−1
0
1
2
Geographic distance (rescaled)

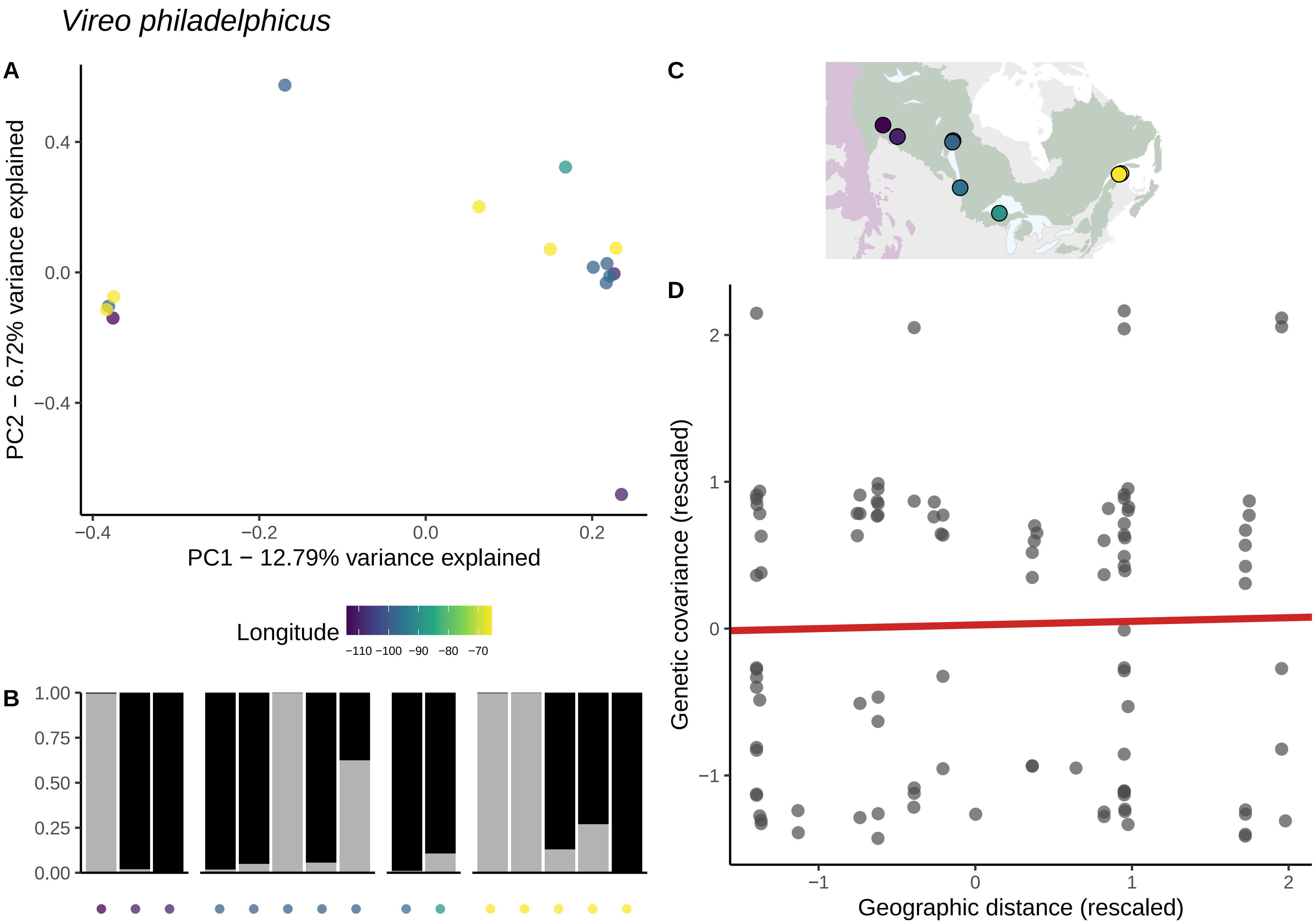
Vireo philadelphicus
A
PC2 − 6.72% variance explained
PC1 − 12.79% variance explained
0.4
0.0
−0.4
−0.2
0.2
Longitude
−110 −100 −90 −80 −70
B
1.00
0.75
0.50
0.25
0.00
C
D
Genetic covariance (rescaled)
Geographic distance (rescaled)
2
1
0
−1

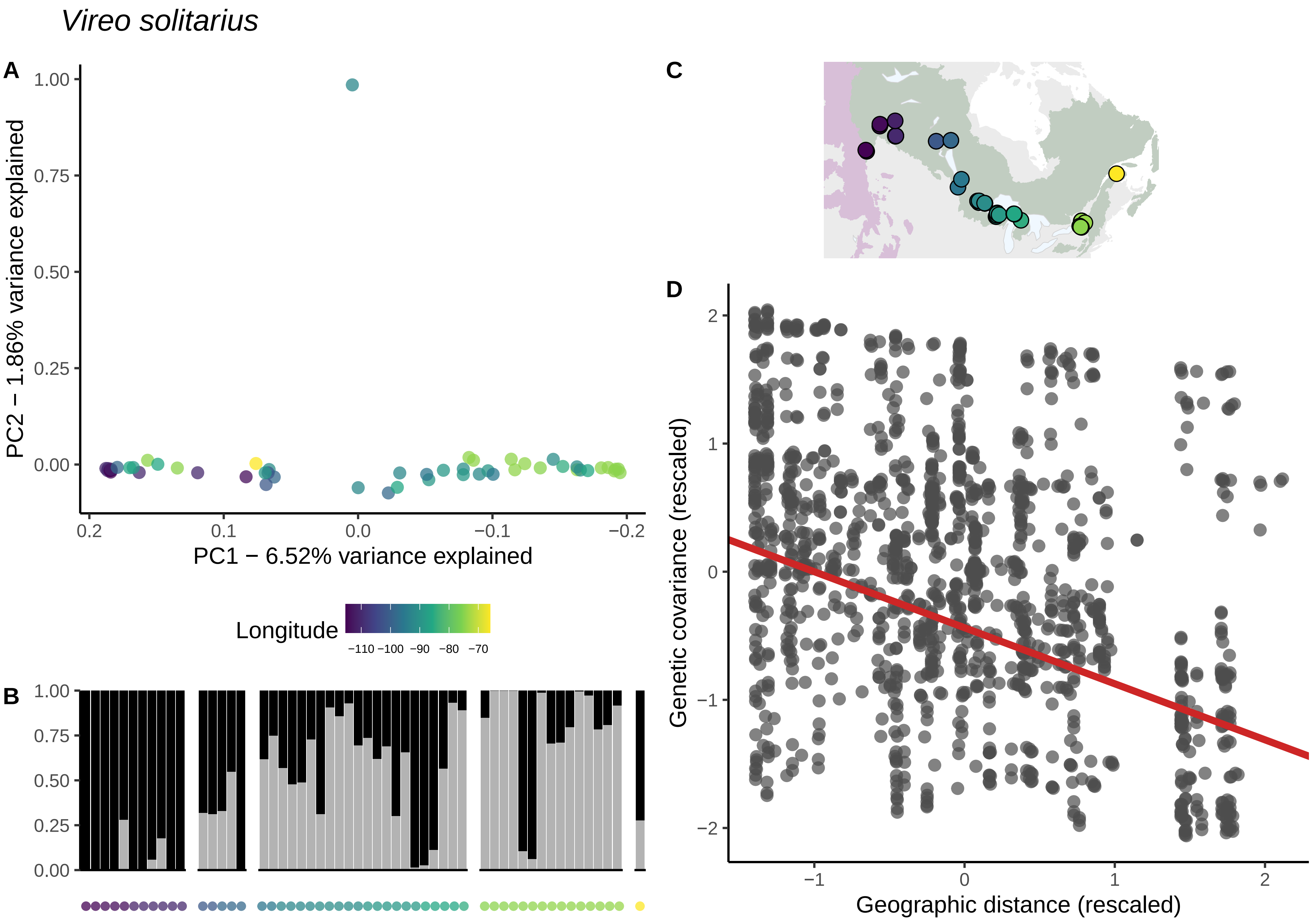

Vireo solitarius
A
B
C
D
PC2 − 1.86% variance explained
PC1 − 6.52% variance explained
Longitude
−110 −100 −90 −80 −70
Genetic covariance (rescaled)
Geographic distance (rescaled)

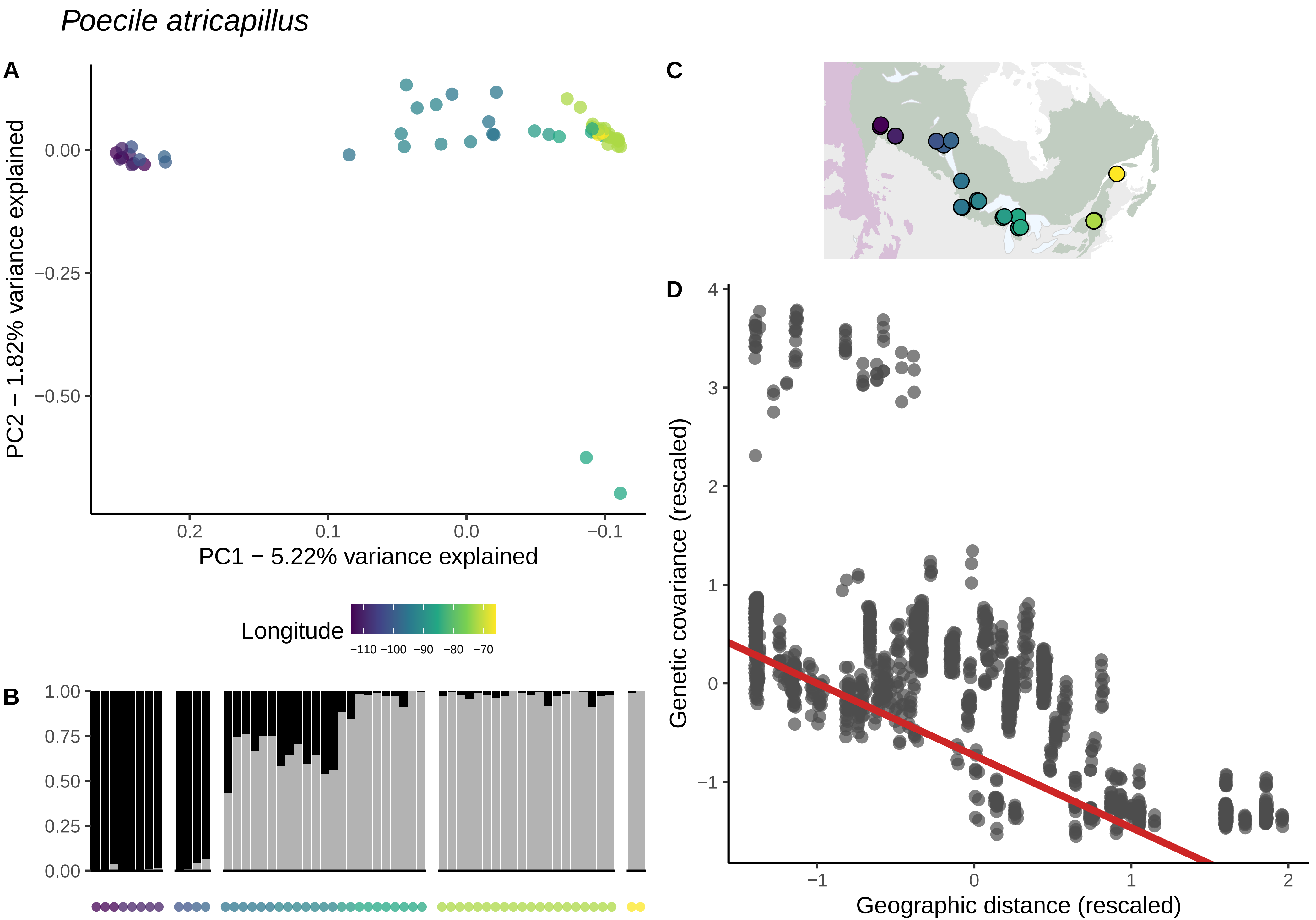
Poecile atricapillus
A
PC2 − 1.82% variance explained
0.00
−0.25
−0.50
0.2
0.1
0.0
−0.1
PC1 − 5.22% variance explained
Longitude
−110 −100 −90 −80 −70
B
1.00
0.75
0.50
0.25
0.00
C
D
Genetic covariance (rescaled)
4
3
2
1
0
−1
−1
0
1
2
Geographic distance (rescaled)

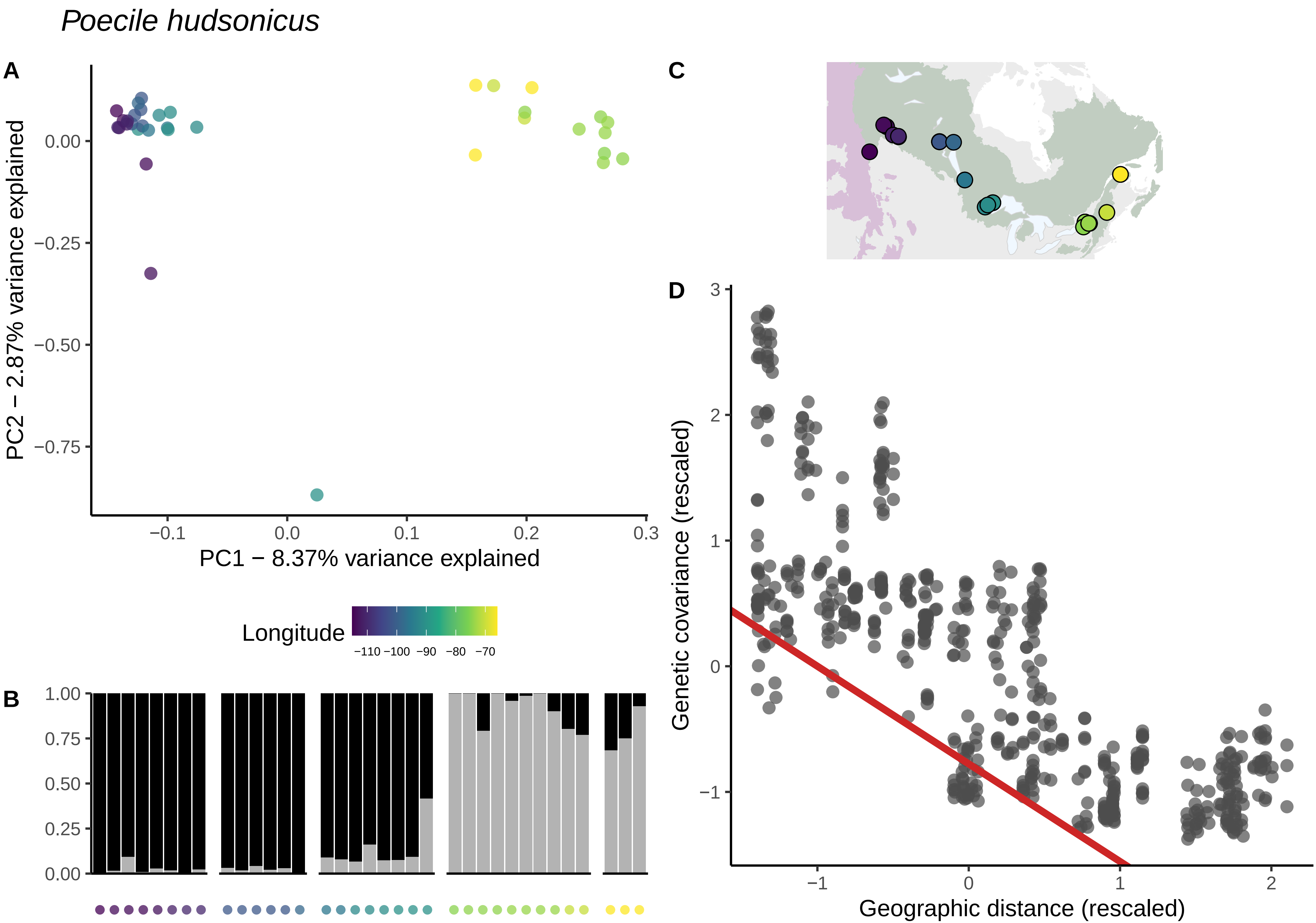
Poecile hudsonicus
A
PC2 − 2.87% variance explained
PC1 − 8.37% variance explained
0.00
−0.25
−0.50
−0.75
−0.1
0.0
0.1
0.2
0.3
Longitude
−110 −100 −90 −80 −70
B
1.00
0.75
0.50
0.25
0.00
C
D
Genetic covariance (rescaled)
Geographic distance (rescaled)
3
2
1
0
−1
2

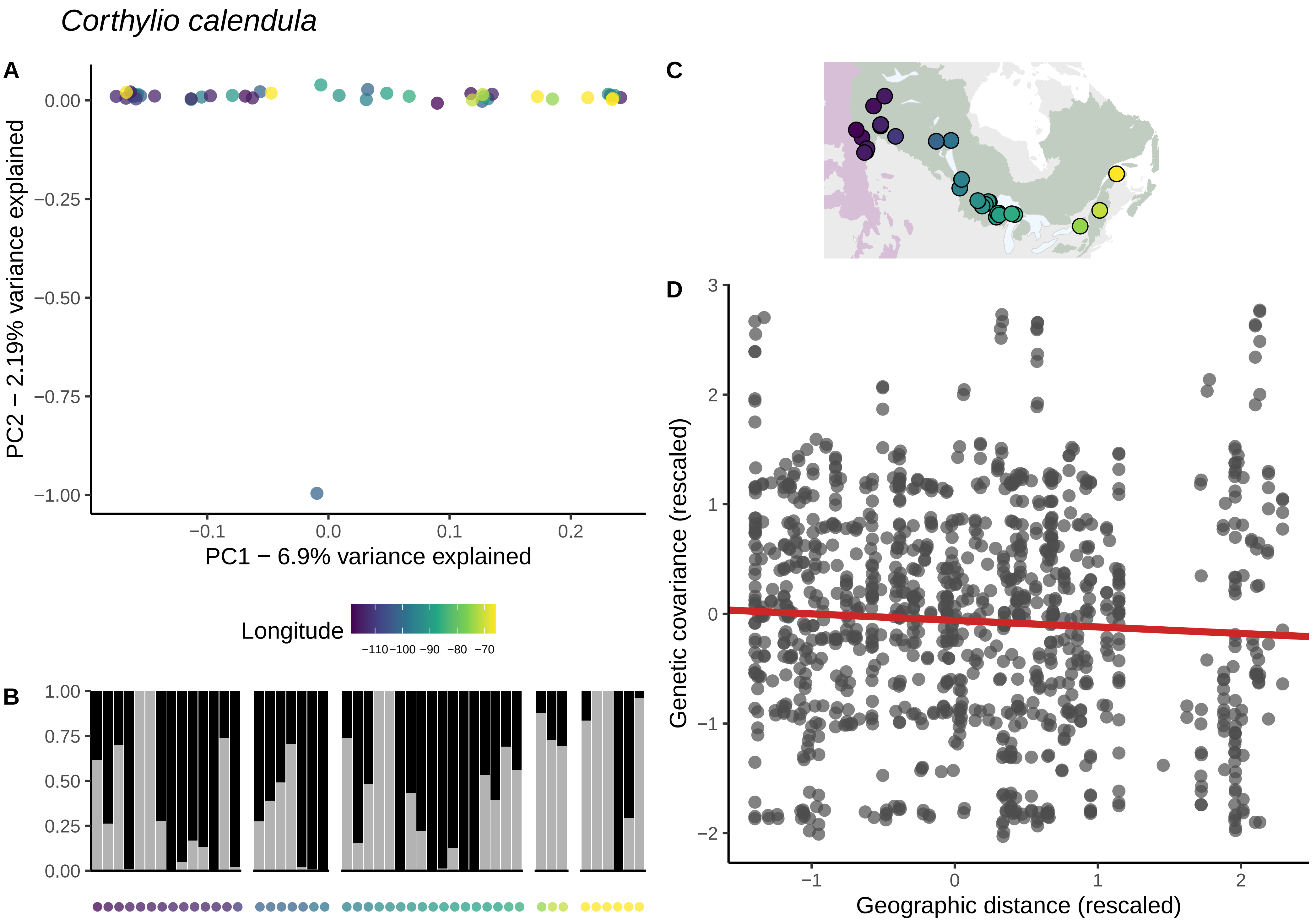

Corthylio calendula
A
PC2 − 2.19% variance explained
0.00
−0.25
−0.50
−0.75
−1.00
−0.1
0.0
0.1
0.2
PC1 − 6.9% variance explained
Longitude
−110−100 −90 −80 −70
B
1.00
0.75
0.50
0.25
0.00
C
D
Genetic covariance (rescaled)
3
2
1
0
−1
−2
Geographic distance (rescaled)

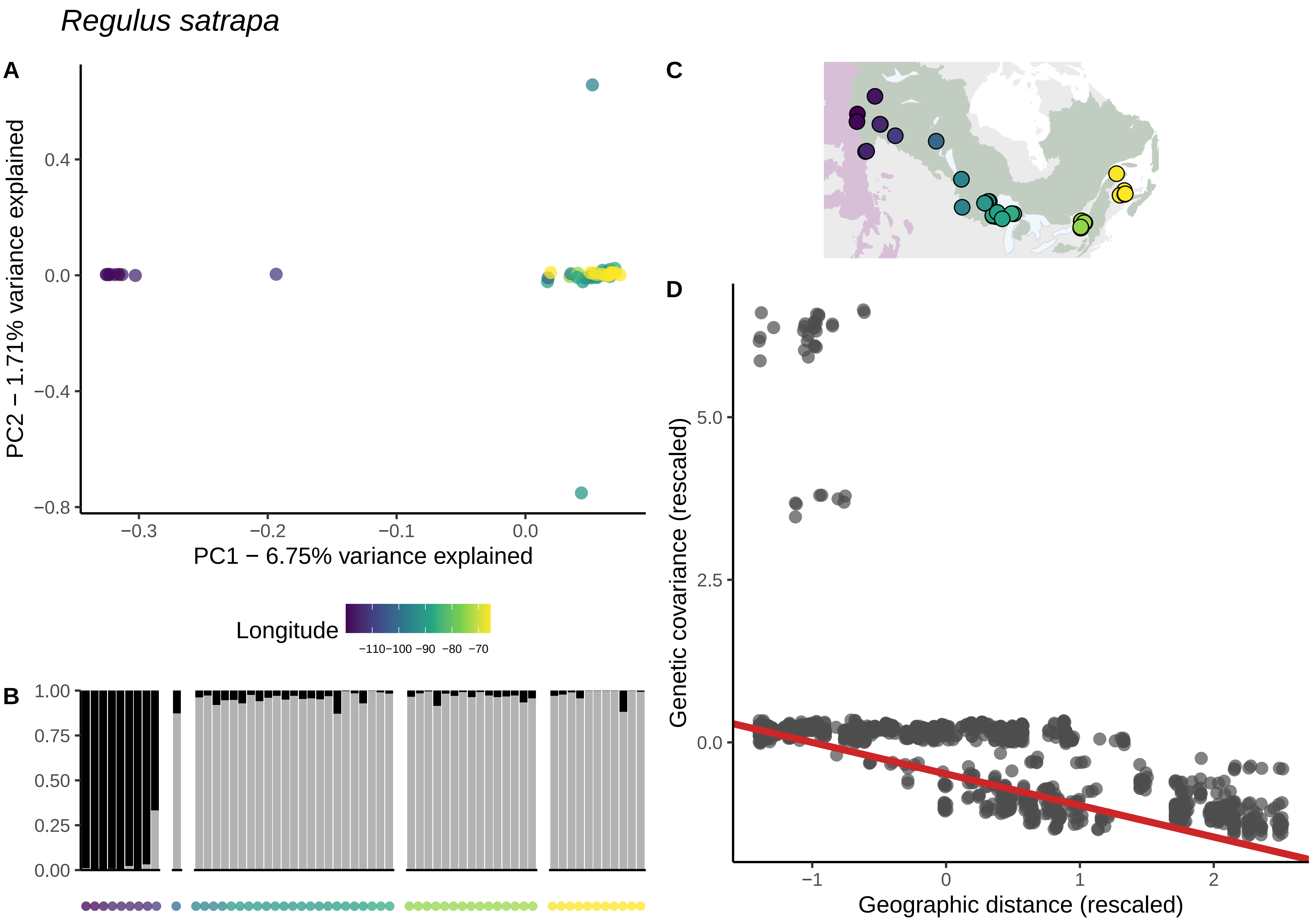

Regulus satrapa
A
PC2 − 1.71% variance explained
0.4
0.0
−0.4
−0.8
−0.3
−0.2
−0.1
0.0
PC1 − 6.75% variance explained
Longitude
−110 −100 −90 −80 −70
B
1.00
0.75
0.50
0.25
0.00
C
D
Genetic covariance (rescaled)
5.0
2.5
0.0
−1
0
1
2
Geographic distance (rescaled)

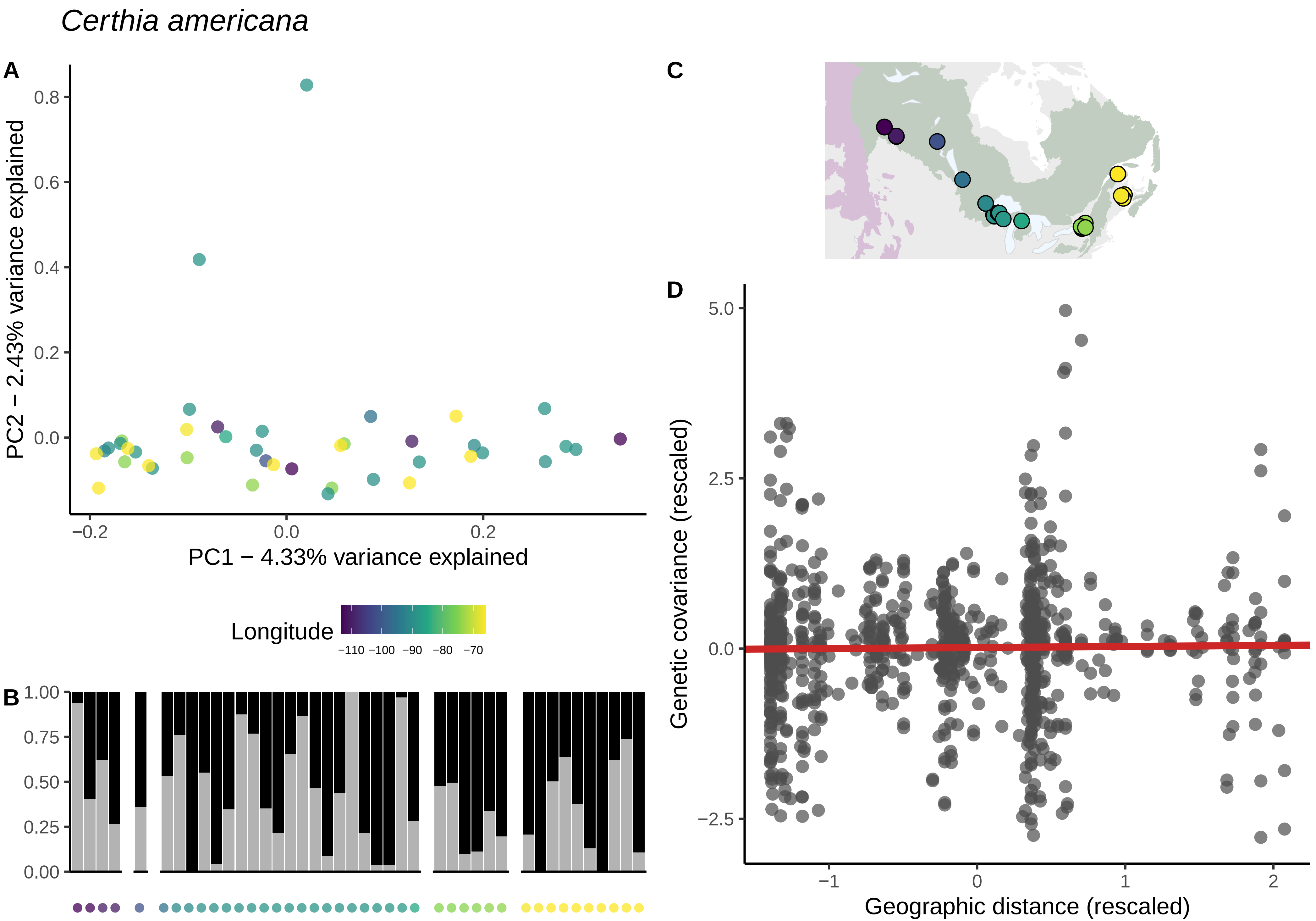

*Certhia americana*
A
B
C
D
PC2 − 2.43% variance explained
PC1 − 4.33% variance explained
Longitude
−110 −100 −90 −80 −70
Genetic covariance (rescaled)
Geographic distance (rescaled)

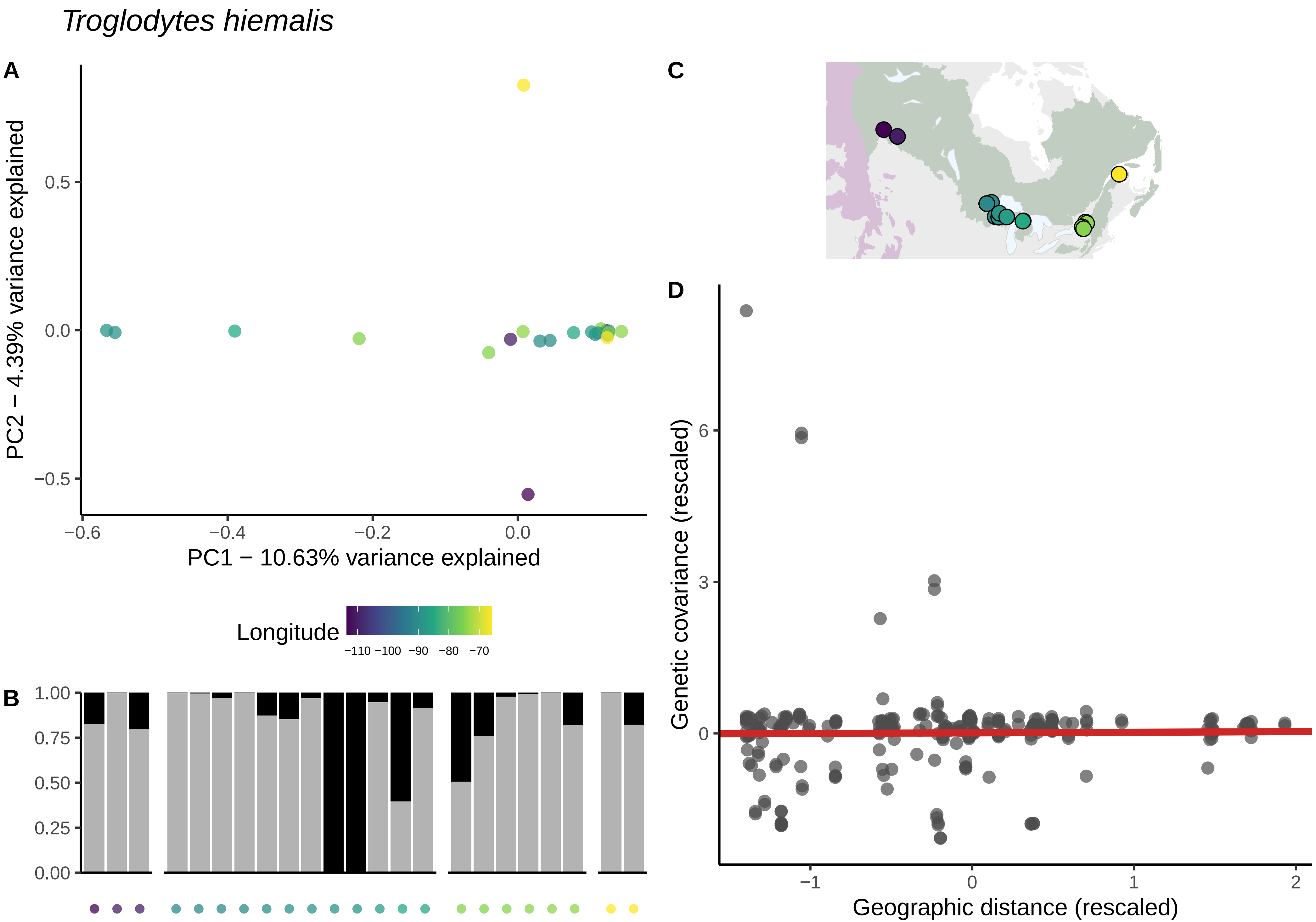
Troglodytes hiemalis
A
PC2 − 4.39% variance explained
PC1 − 10.63% variance explained
0.5
0.0
−0.5
−0.6
−0.4
−0.2
0.0
Longitude
−110 −100 −90 −80 −70
B
1.00
0.75
0.50
0.25
0.00
C
D
Genetic covariance (rescaled)
Geographic distance (rescaled)
6
3
0
−1
0
1
2

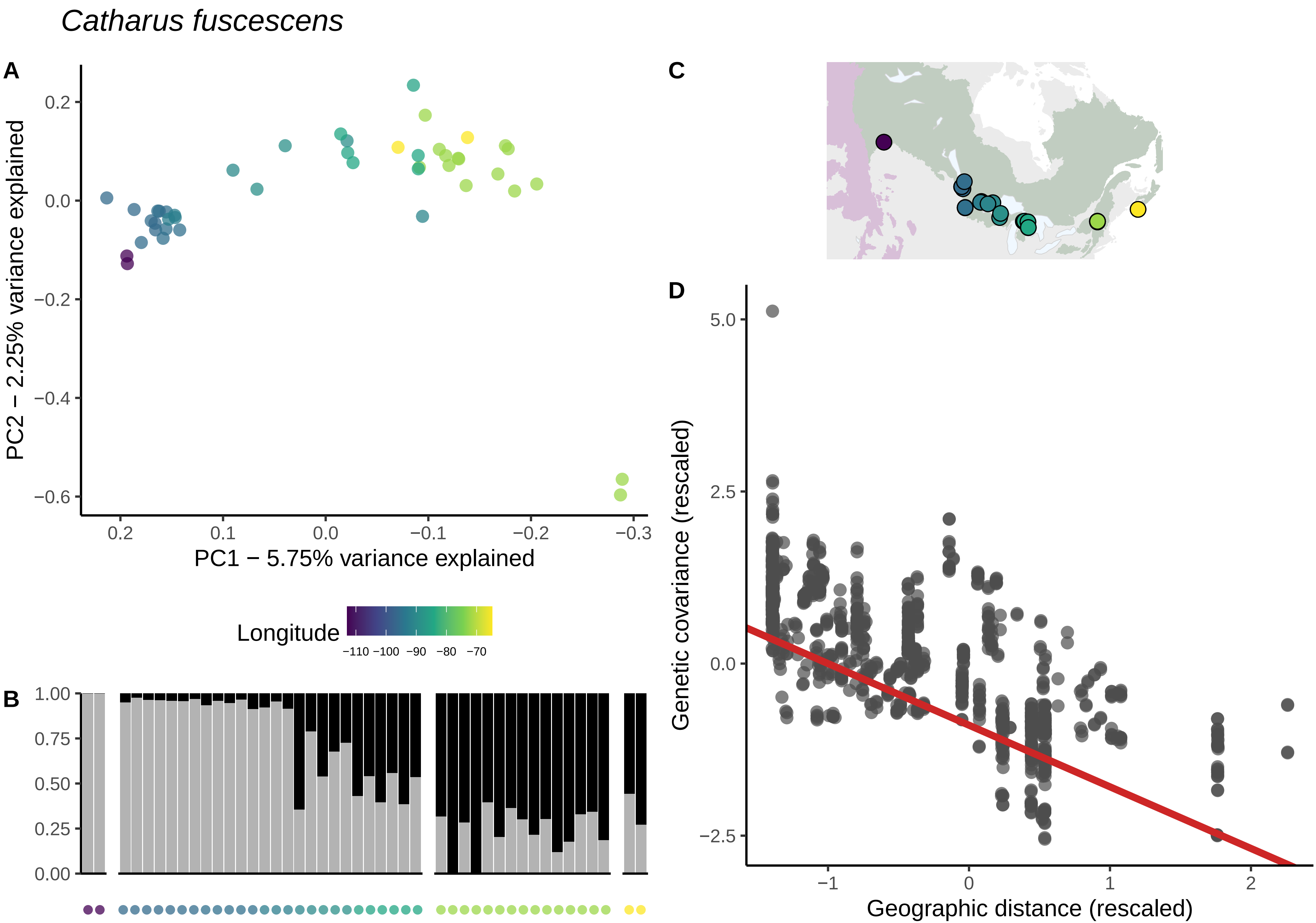

Catharus fuscescens
A
PC2 − 2.25% variance explained
PC1 − 5.75% variance explained
0.2
0.0
−0.2
−0.4
−0.6
0.2
0.1
0.0
−0.1
−0.2
−0.3
Longitude
−110 −100 −90 −80 −70
B
1.00
0.75
0.50
0.25
0.00
C
D
Genetic covariance (rescaled)
Geographic distance (rescaled)
5.0
2.5
0.0
−2.5
−1
0
1
2

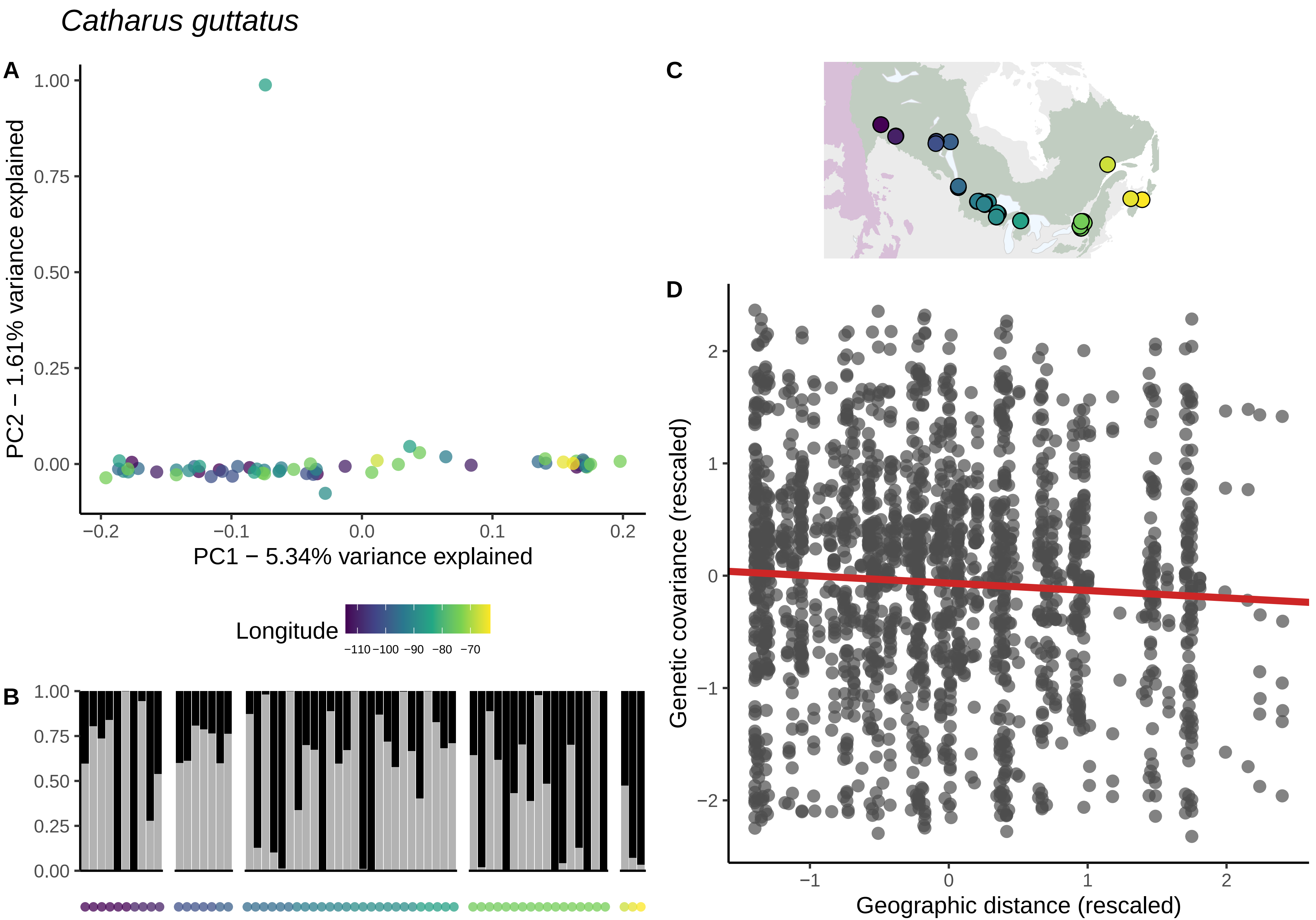

Catharus guttatus
A
B
C
D
1.00
0.75
0.50
0.25
0.00
PC2 − 1.61% variance explained
PC1 − 5.34% variance explained
−0.2
−0.1
0.0
0.1
0.2
Longitude
−110 −100 −90 −80 −70
Genetic covariance (rescaled)
Geographic distance (rescaled)
2
1
0
−1
−2

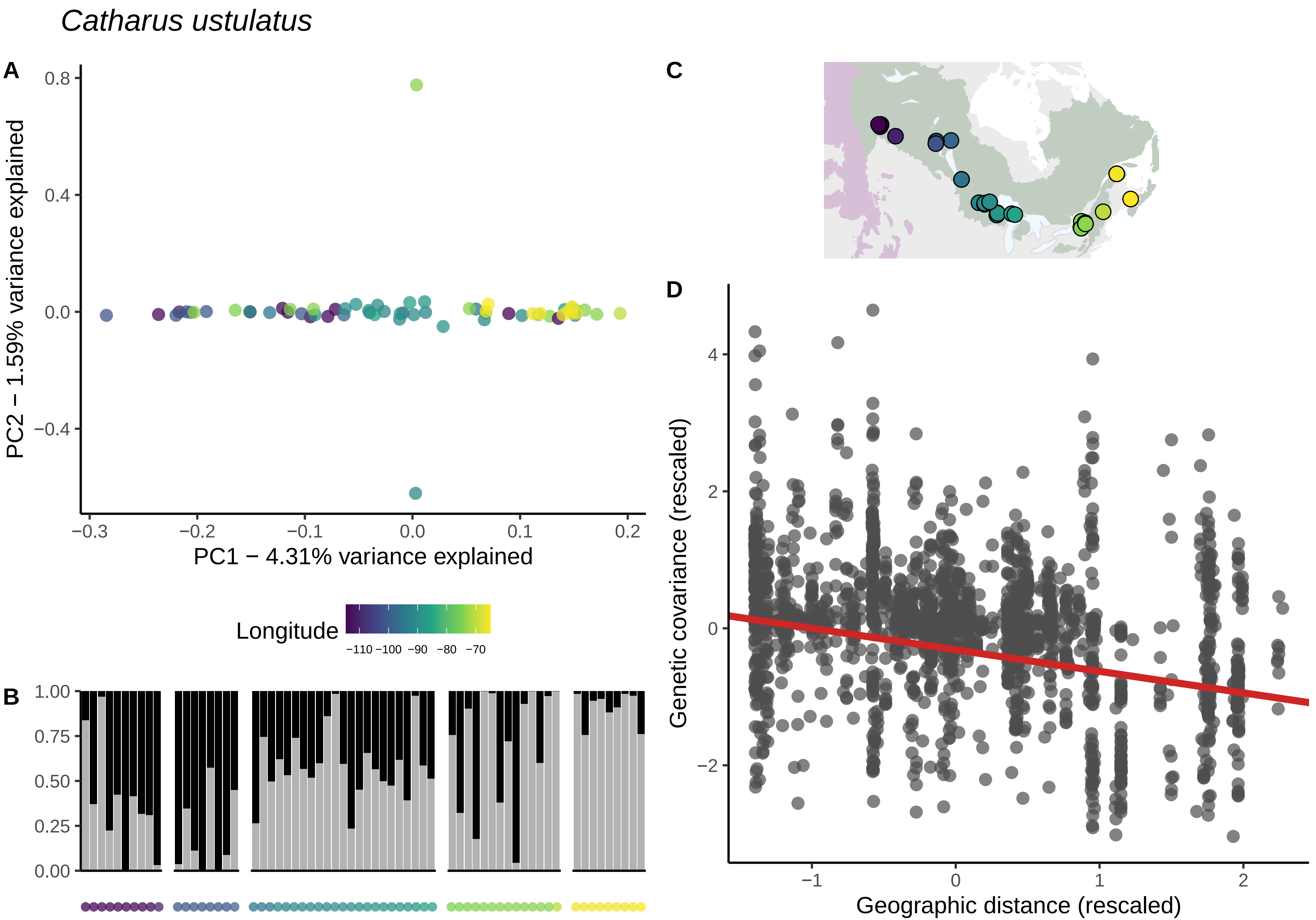

*Catharus ustulatus*
A
PC2 − 1.59% variance explained
PC1 − 4.31% variance explained
0.8
0.4
0.0
−0.4
−0.3
−0.2
−0.1
0.0
0.1
0.2
Longitude
−110 −100 −90 −80 −70
B
1.00
0.75
0.50
0.25
0.00
C
D
Genetic covariance (rescaled)
Geographic distance (rescaled)
4
2
0
−2
−1
0
1
2

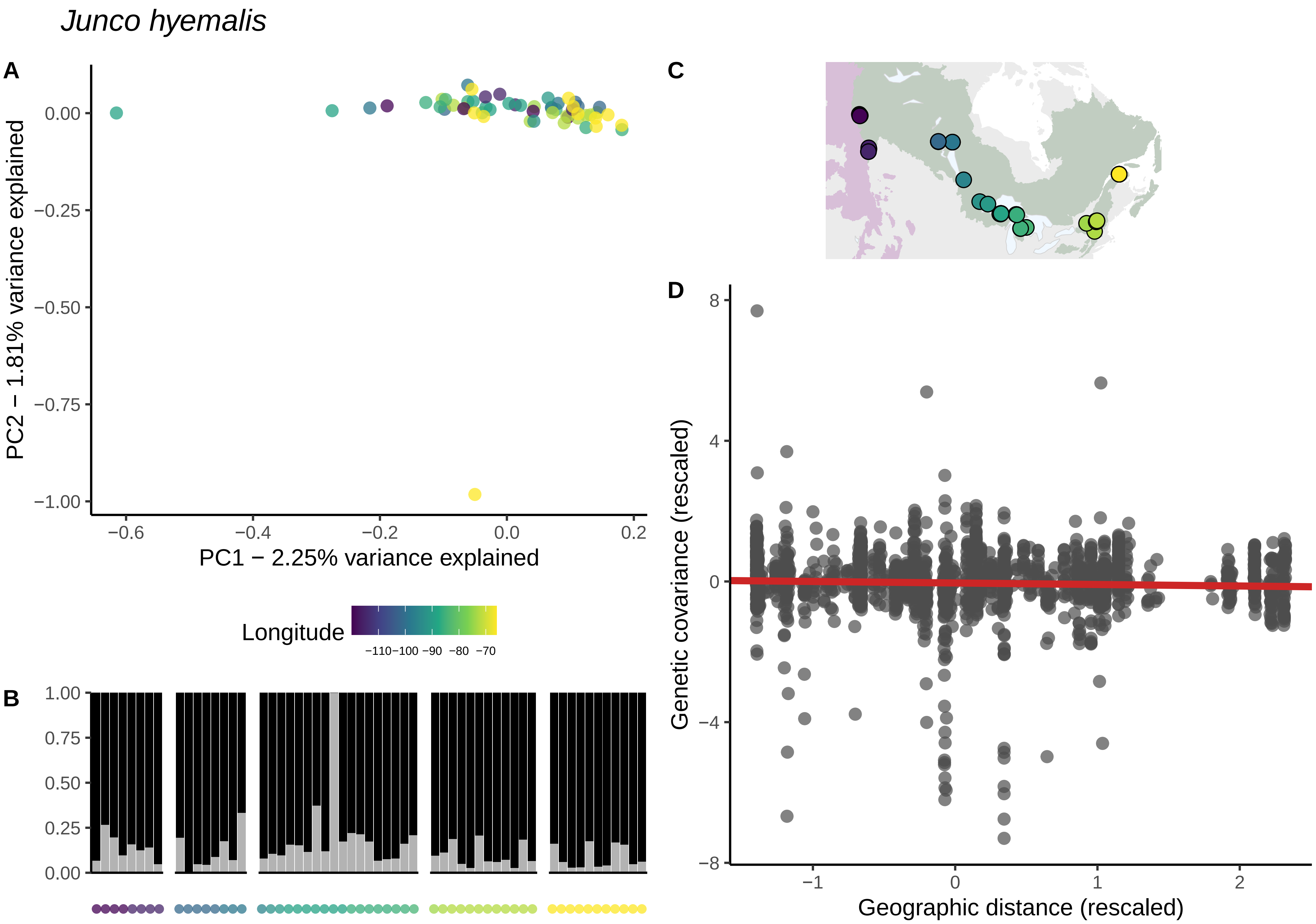
Junco hyemalis
A
PC2 − 1.81% variance explained
0.00
−0.25
−0.50
−0.75
−1.00
−0.6
−0.4
−0.2
0.0
0.2
PC1 − 2.25% variance explained
Longitude
−110−100 −90 −80 −70
B
1.00
0.75
0.50
0.25
0.00
C
D
Genetic covariance (rescaled)
8
4
0
−4
−8
−1
0
1
2
Geographic distance (rescaled)

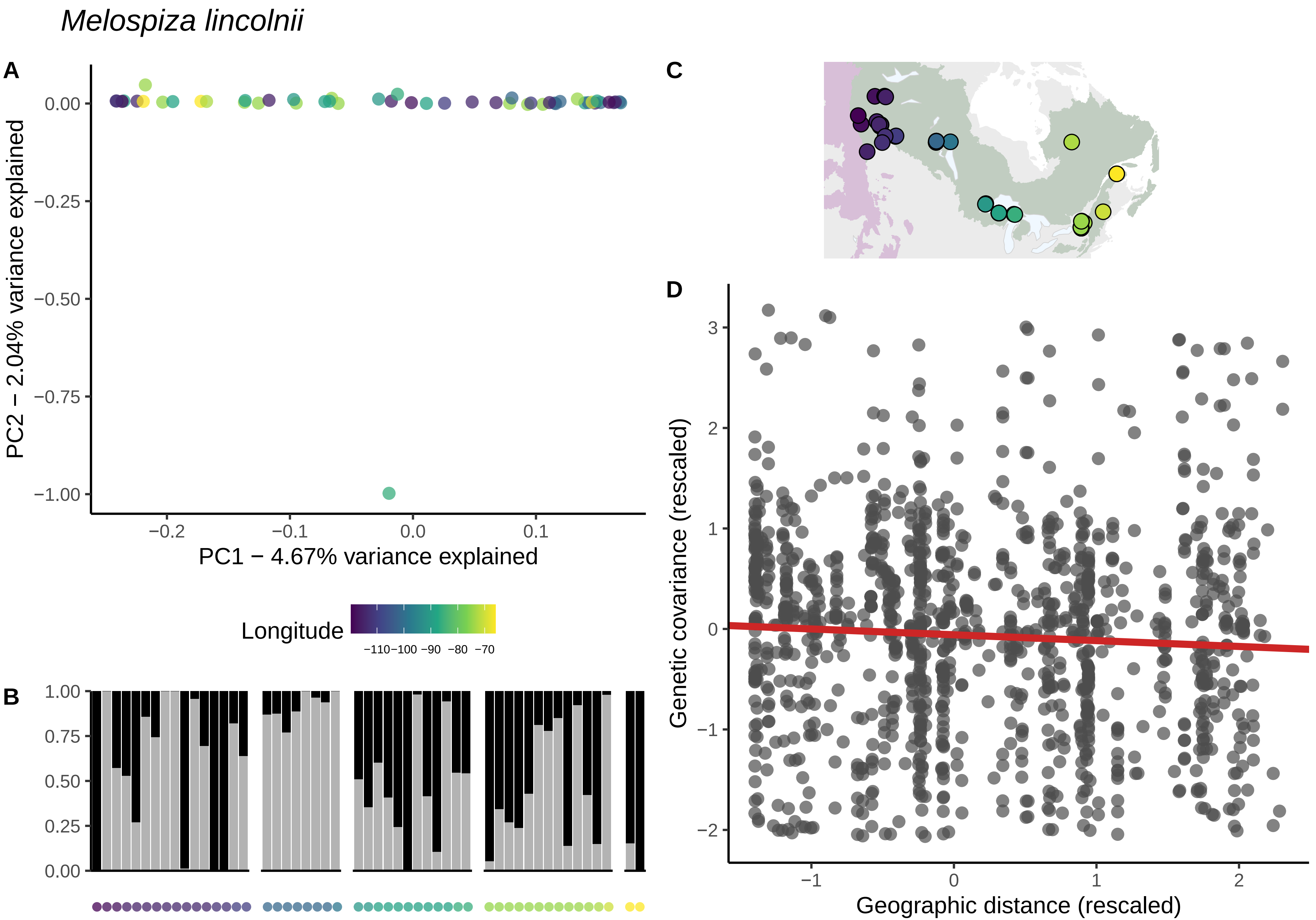

Melospiza lincolnii
A
B
C
D
PC2 − 2.04% variance explained
PC1 − 4.67% variance explained
0.00
−0.25
−0.50
−0.75
−1.00
−0.2
−0.1
0.0
0.1
Longitude
−110−100 −90 −80 −70
1.00
0.75
0.50
0.25
0.00
Genetic covariance (rescaled)
Geographic distance (rescaled)
3
2
1
0
−1
−2
−1
0
1
2

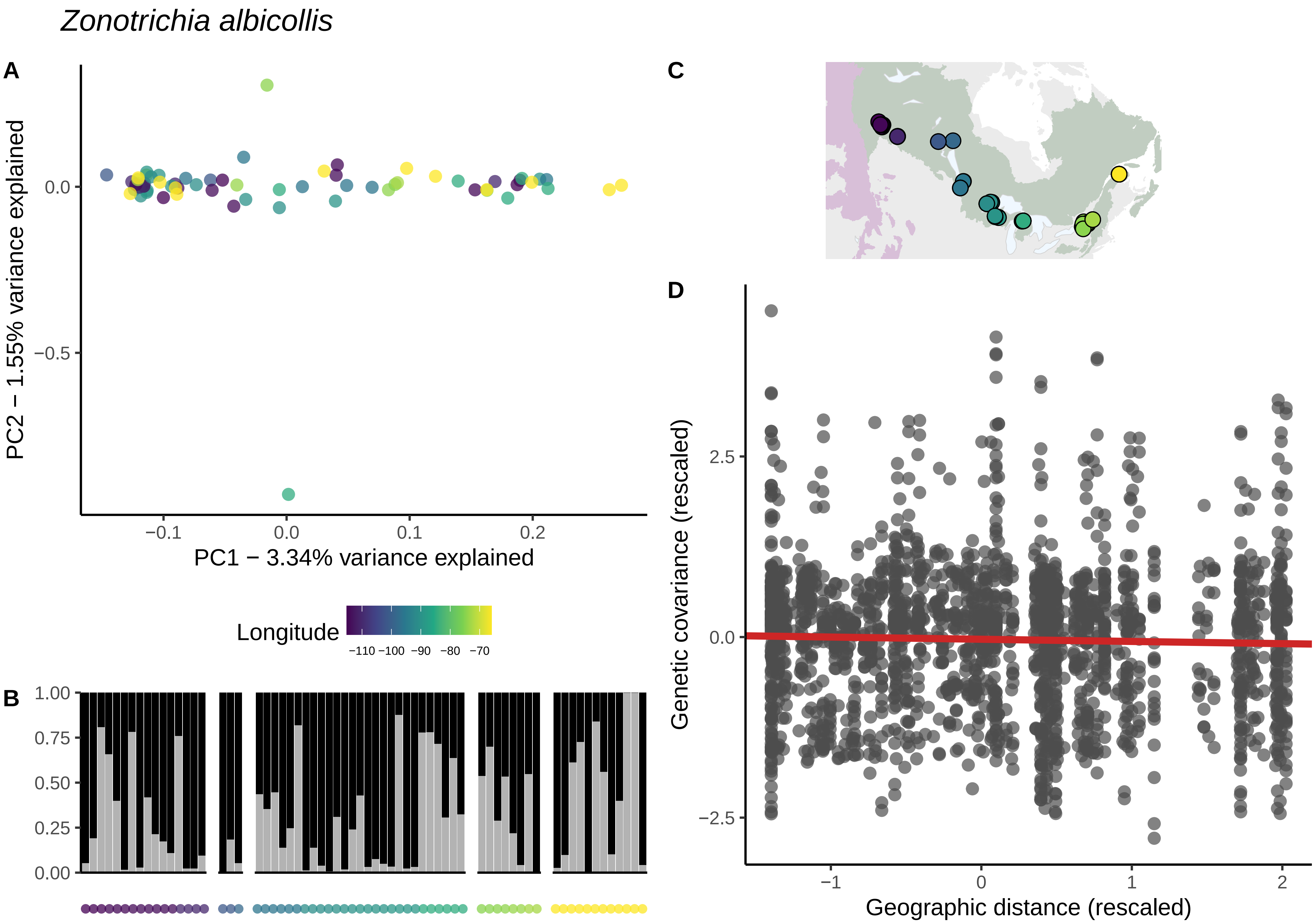

Zonotrichia albicollis
A
C
D
B
0.0
−0.5
PC2 − 1.55% variance explained
−0.1
0.0
0.1
0.2
PC1 − 3.34% variance explained
Longitude
−110 −100 −90 −80 −70
1.00
0.75
0.50
0.25
0.00
2.5
0.0
−2.5
Genetic covariance (rescaled)
−1
0
1
2
Geographic distance (rescaled)

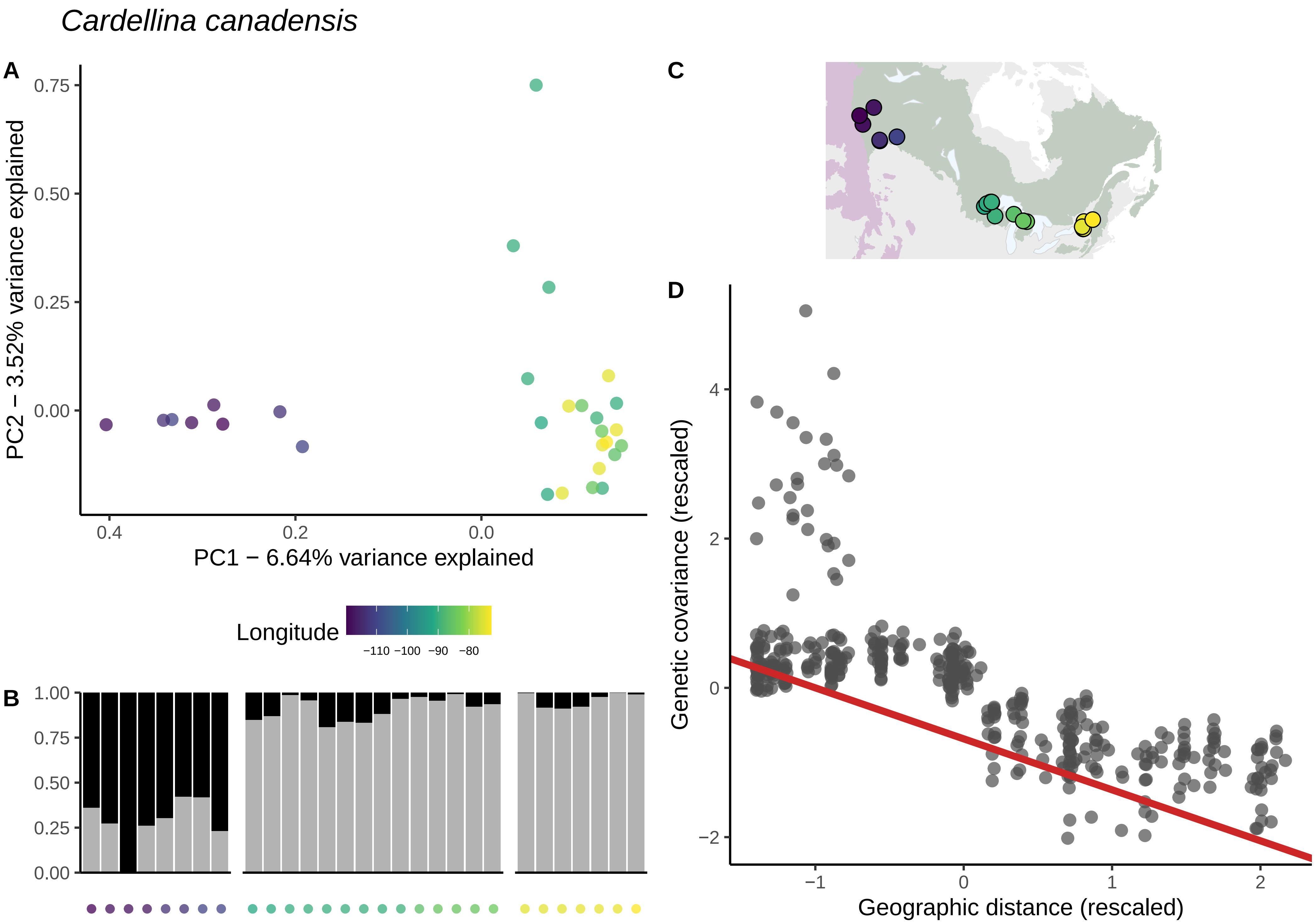

Cardellina canadensis
A
C
D
B
PC2 − 3.52% variance explained
PC1 − 6.64% variance explained
0.75
0.50
0.25
0.00
0.4
0.2
0.0
Longitude
−110 −100 −90 −80
1.00
0.75
0.50
0.25
0.00
Genetic covariance (rescaled)
Geographic distance (rescaled)
4
2
0
−2
−1
0
1
2

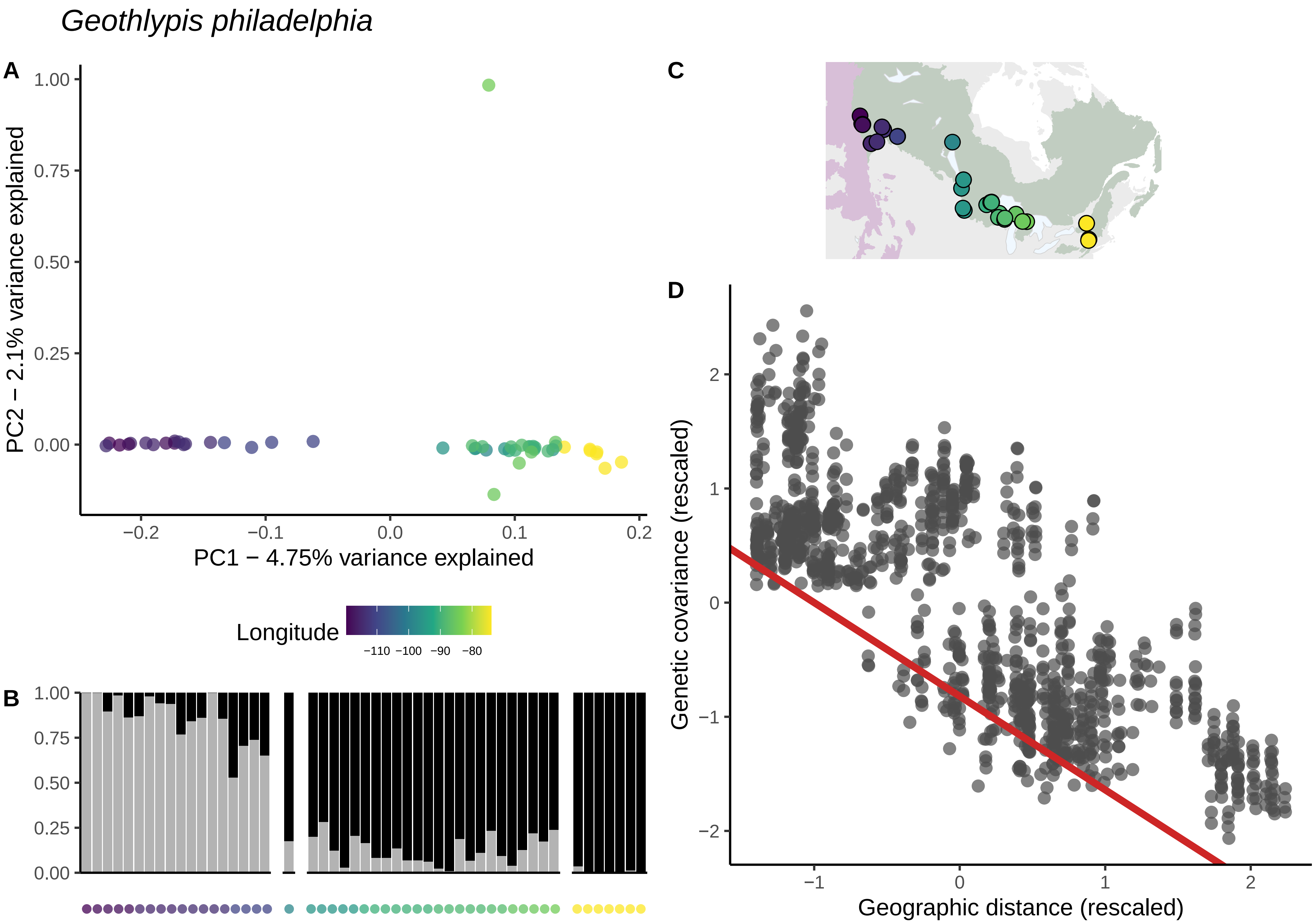
Geothlypis philadelphia
A
B
C
D
PC2 − 2.1% variance explained
PC1 − 4.75% variance explained
1.00
0.75
0.50
0.25
0.00
−0.2
−0.1
0.0
0.1
0.2
Longitude
−110 −100 −90 −80
Genetic covariance (rescaled)
Geographic distance (rescaled)
2
1
0
−1
−2

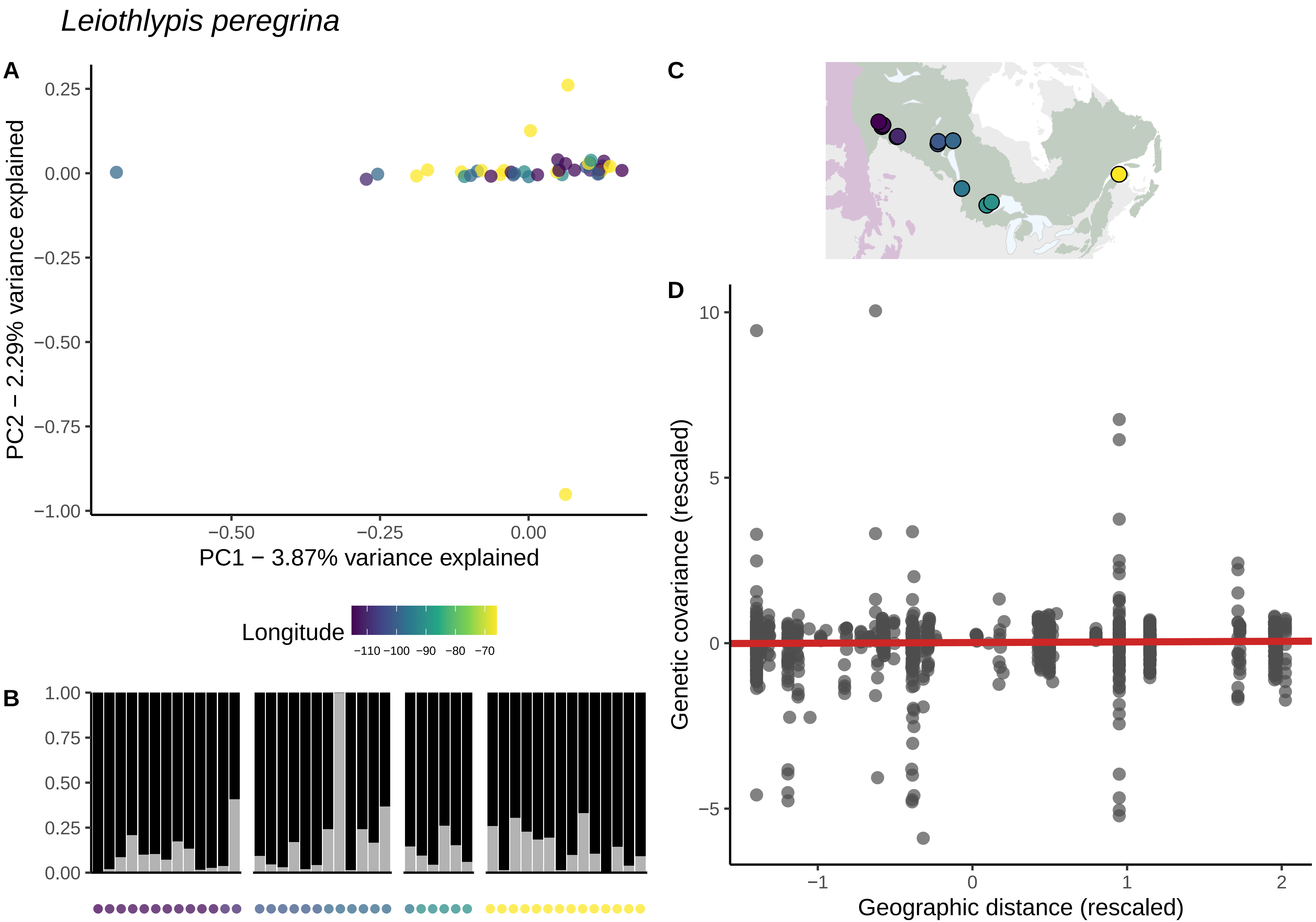
Leiothlypis peregrina
A
B
C
D
PC1 − 3.87% variance explained
PC2 − 2.29% variance explained
Longitude
−110 −100 −90 −80 −70
Genetic covariance (rescaled)
Geographic distance (rescaled)

*Leiothlypis ruficapilla*

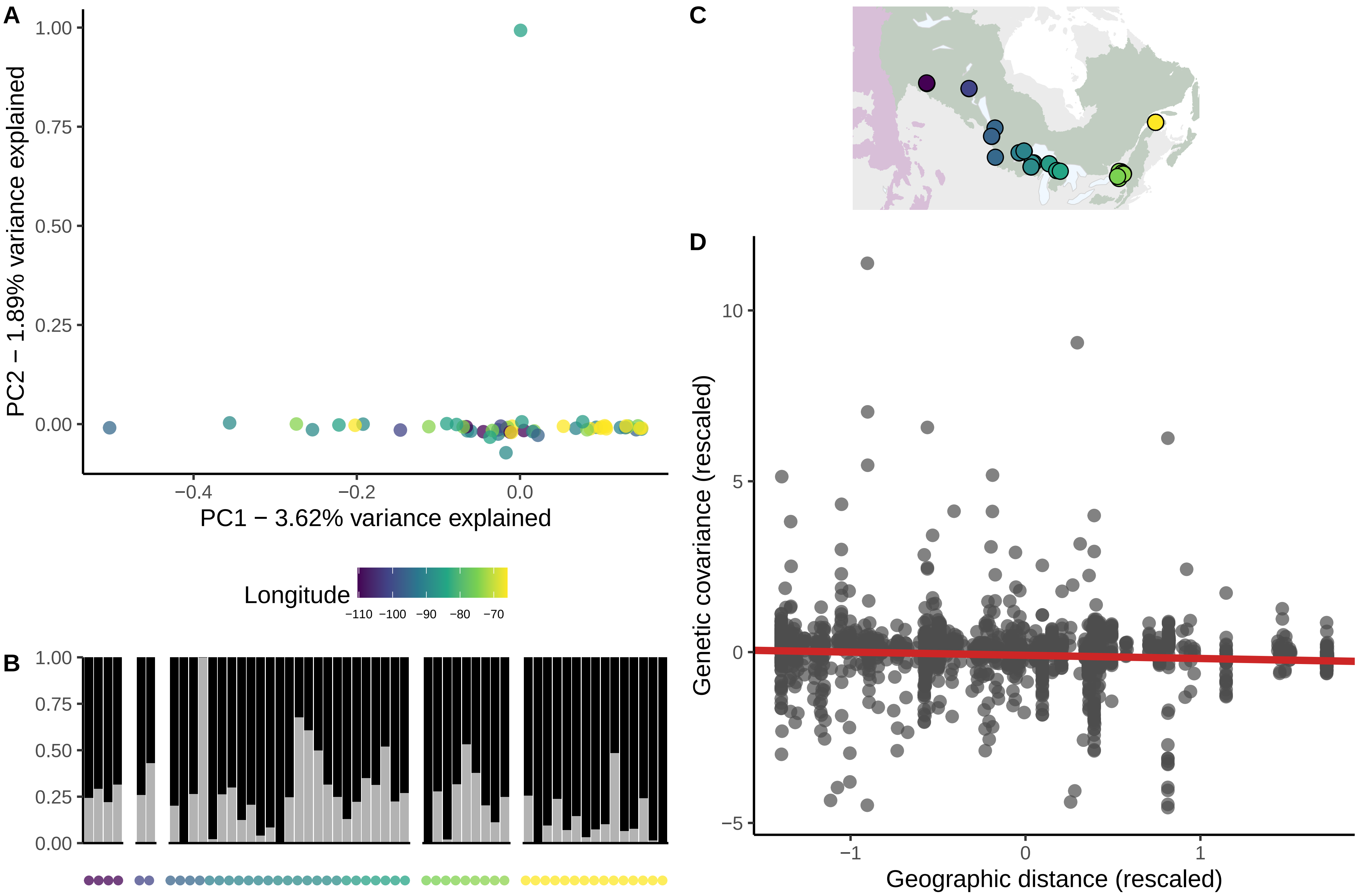

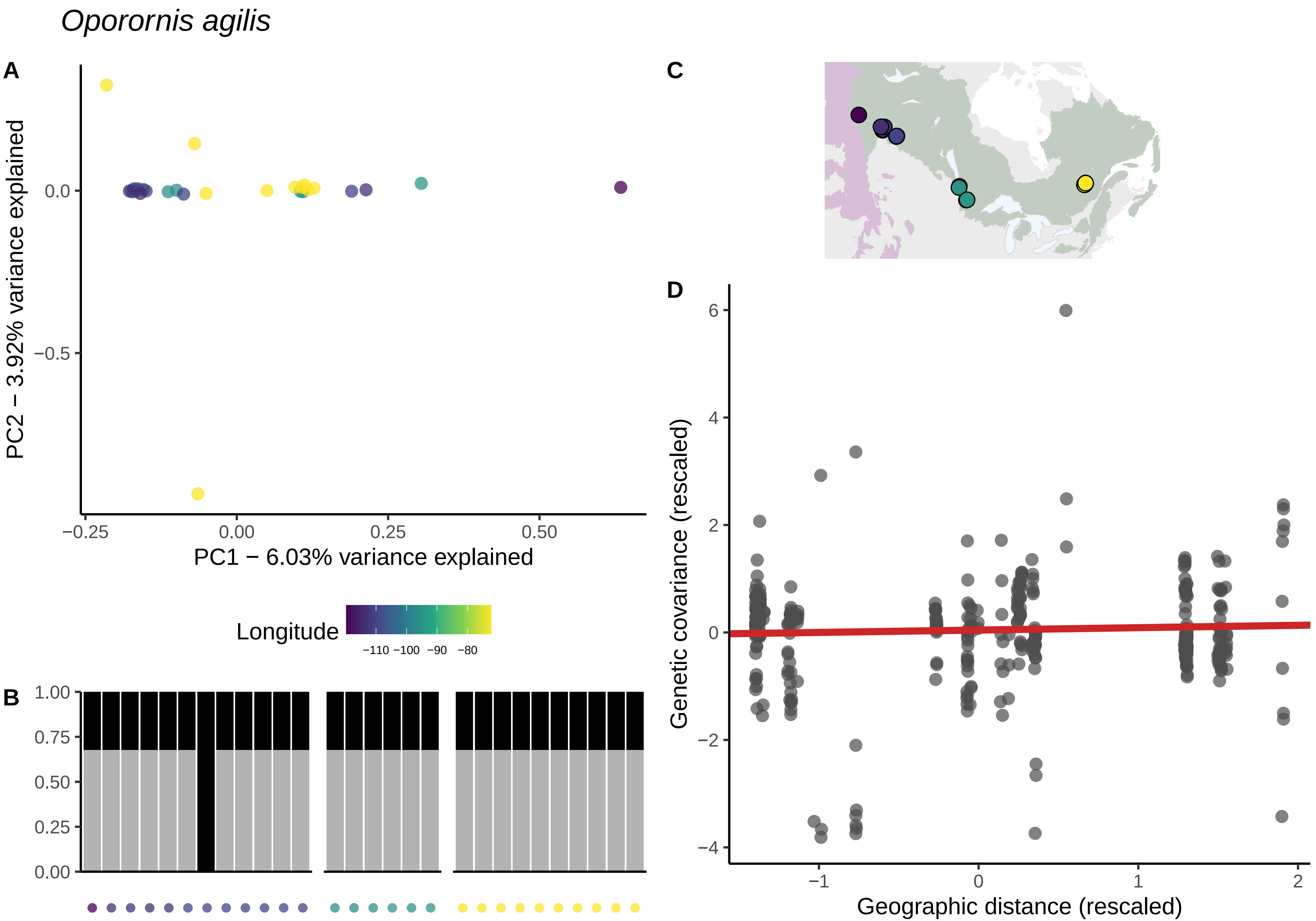

Oporornis agilis
A
PC2 − 3.92% variance explained
0.0
−0.5
−0.25
0.00
0.25
0.50
PC1 − 6.03% variance explained
Longitude
−110 −100 −90 −80
B
1.00
0.75
0.50
0.25
0.00
C
D
6
4
2
0
−2
−4
Genetic covariance (rescaled)
−1
0
1
2
Geographic distance (rescaled)

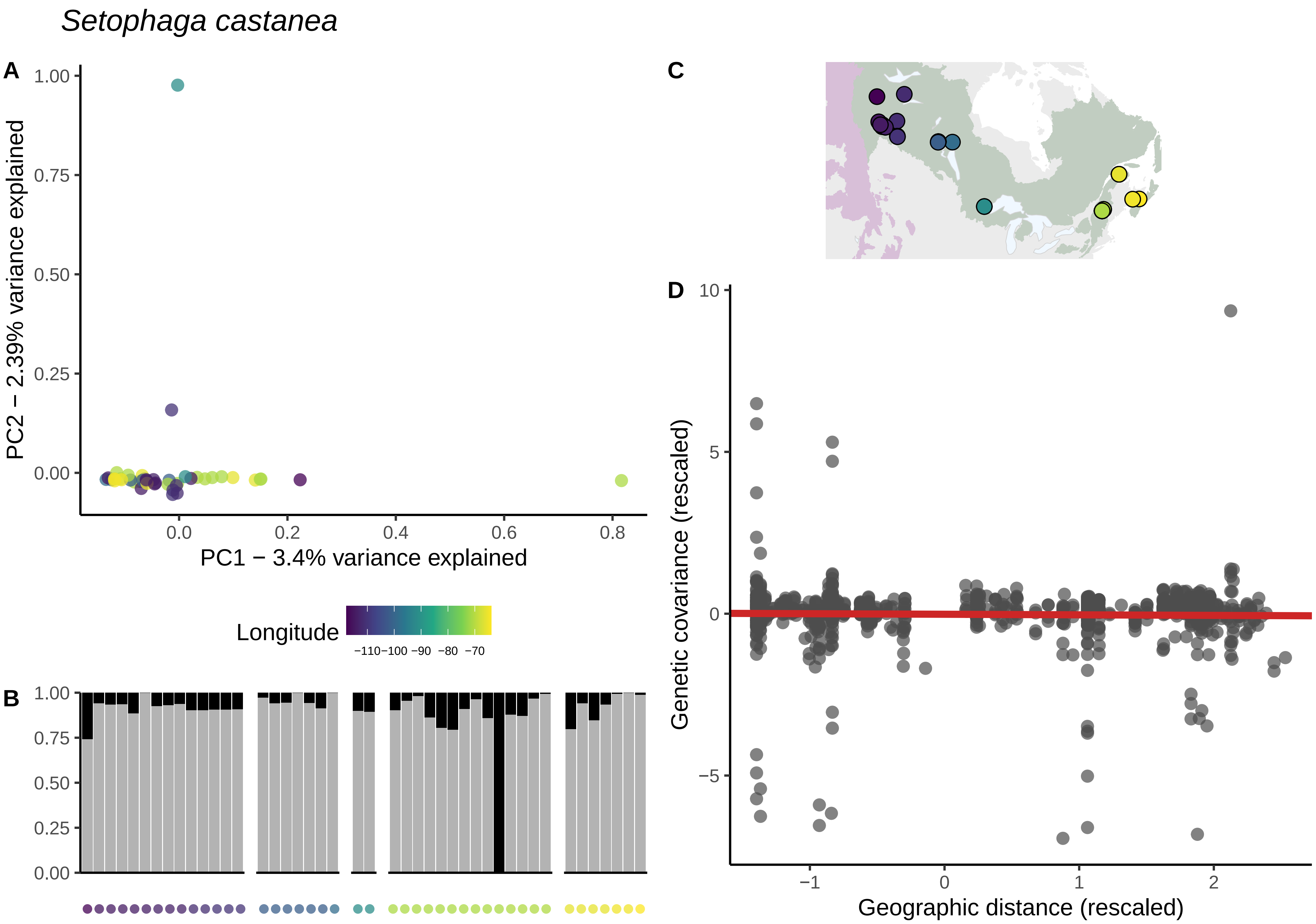

Setophaga castanea
A
PC2 − 2.39% variance explained
PC1 − 3.4% variance explained
Longitude
−110−100 −90 −80 −70
B
C
D
Genetic covariance (rescaled)
Geographic distance (rescaled)

*Setophaga coronata*

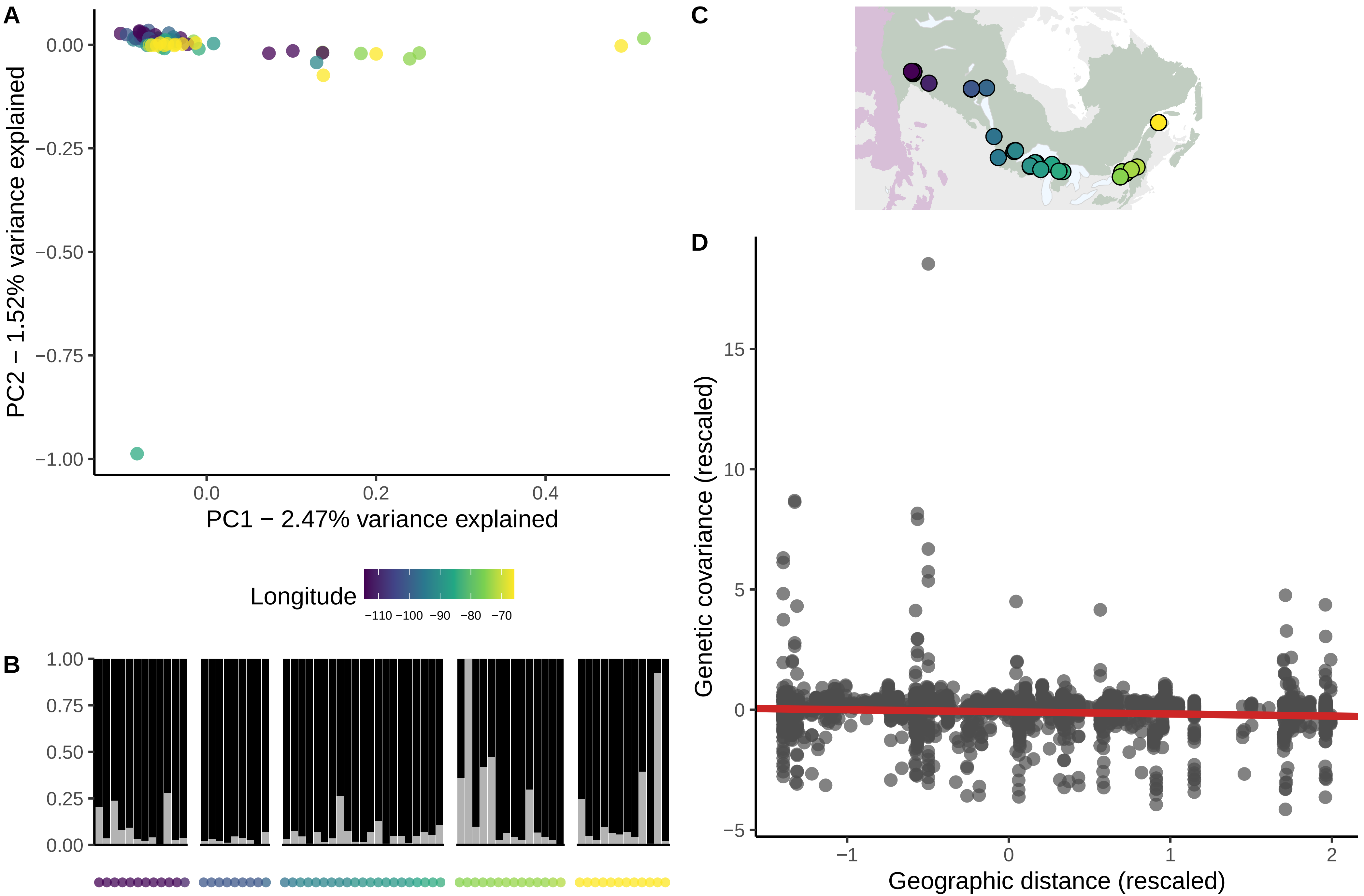

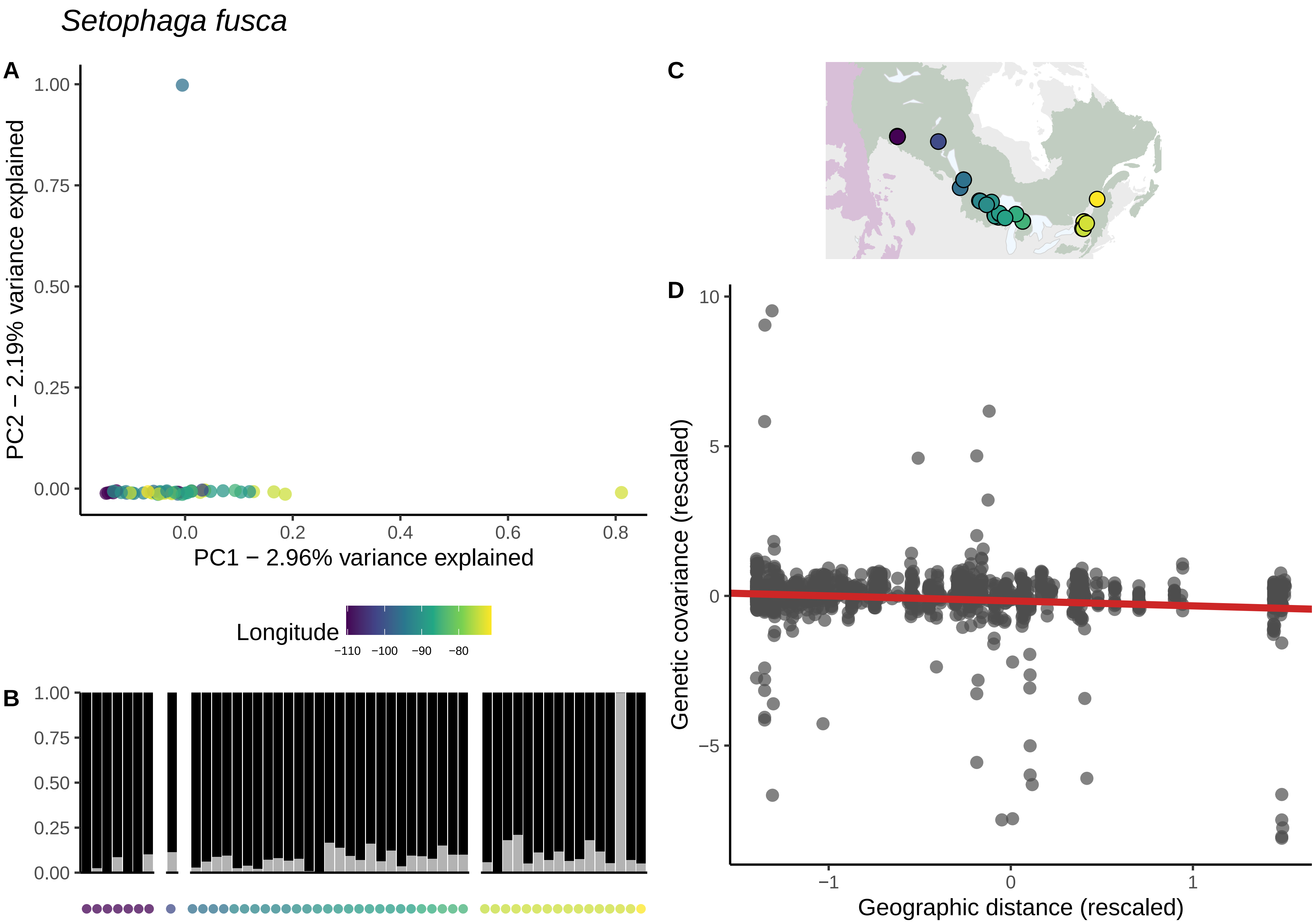

Setophaga fusca
A
B
C
D
PC2 − 2.19% variance explained
PC1 − 2.96% variance explained
1.00
0.75
0.50
0.25
0.00
0.0
0.2
0.4
0.6
0.8
Longitude
−110
−100
−90
−80
Genetic covariance (rescaled)
Geographic distance (rescaled)
10
5
0
−5
−1
1

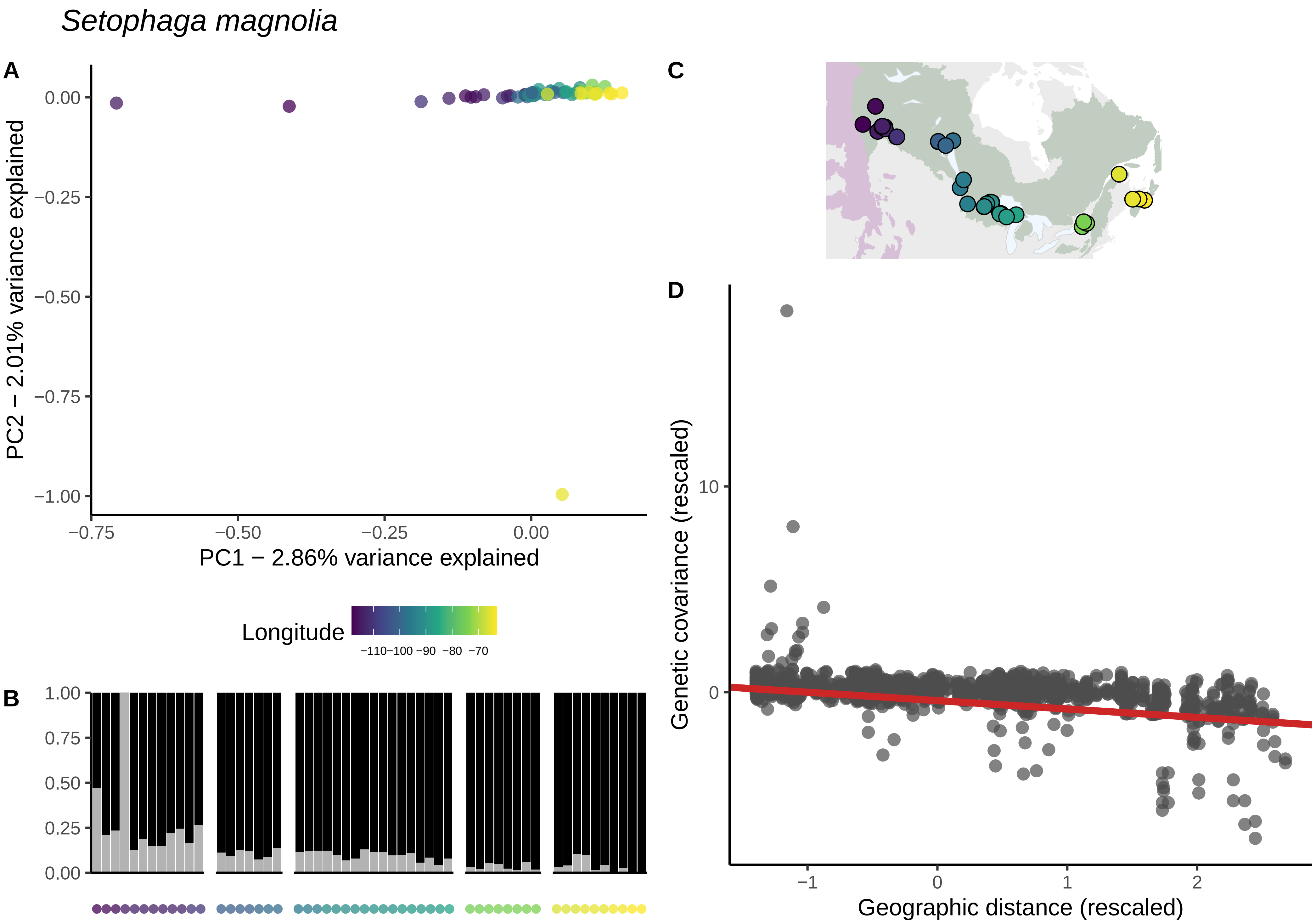

Setophaga magnolia
A
PC2 − 2.01% variance explained
0.00
−0.25
−0.50
−0.75
−1.00
−0.75
−0.50
−0.25
0.00
PC1 − 2.86% variance explained
Longitude
−110−100 −90 −80 −70
B
1.00
0.75
0.50
0.25
0.00
C
D
Genetic covariance (rescaled)
10
0
−1
0
1
2
Geographic distance (rescaled)

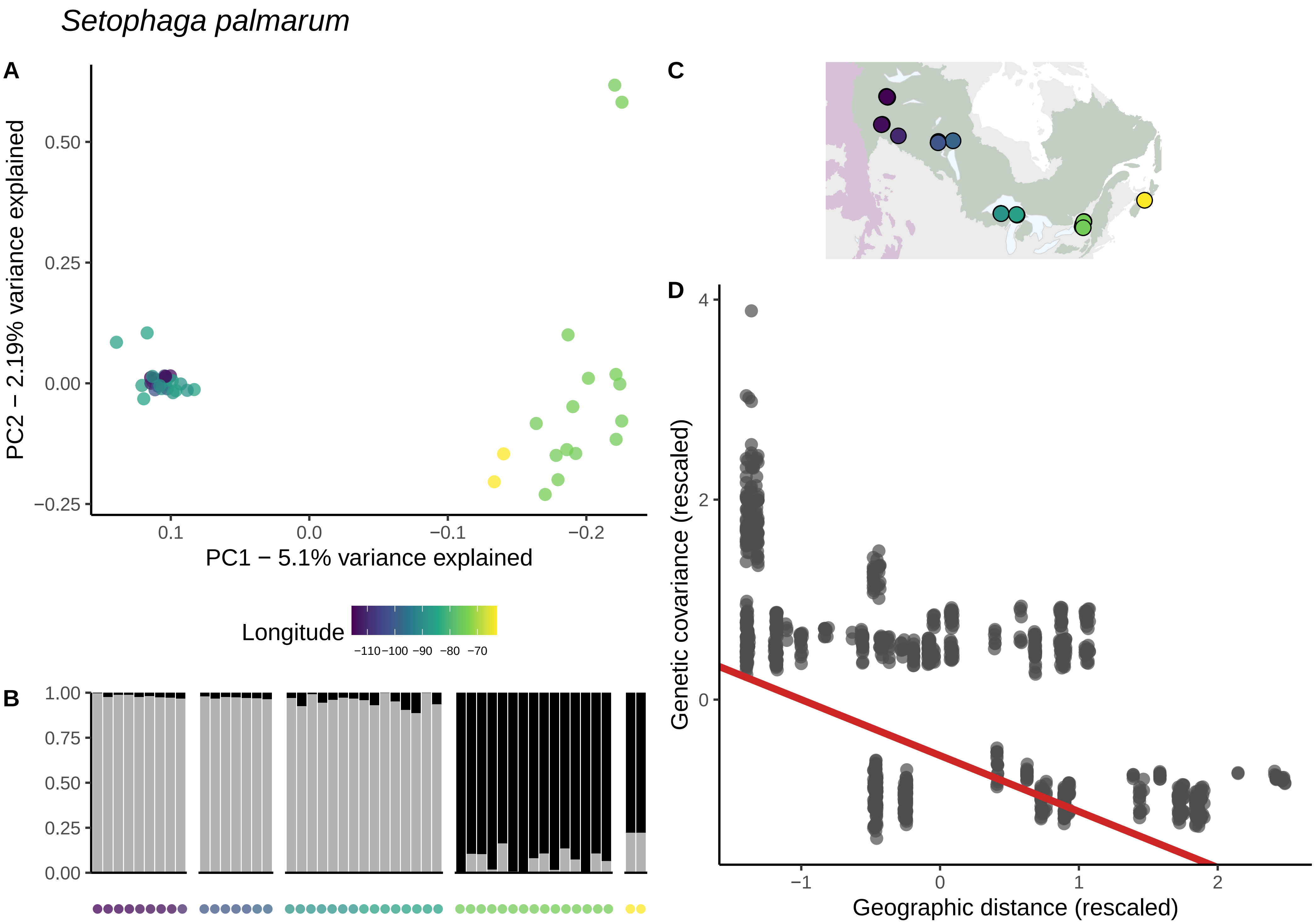
Setophaga palmarum
A
PC2 − 2.19% variance explained
PC1 − 5.1% variance explained
0.50
0.25
0.00
−0.25
0.1
0.0
−0.1
−0.2
Longitude
−110−100 −90 −80 −70
B
1.00
0.75
0.50
0.25
0.00
C
D
Genetic covariance (rescaled)
Geographic distance (rescaled)
4
2
0
−1
0
1
2

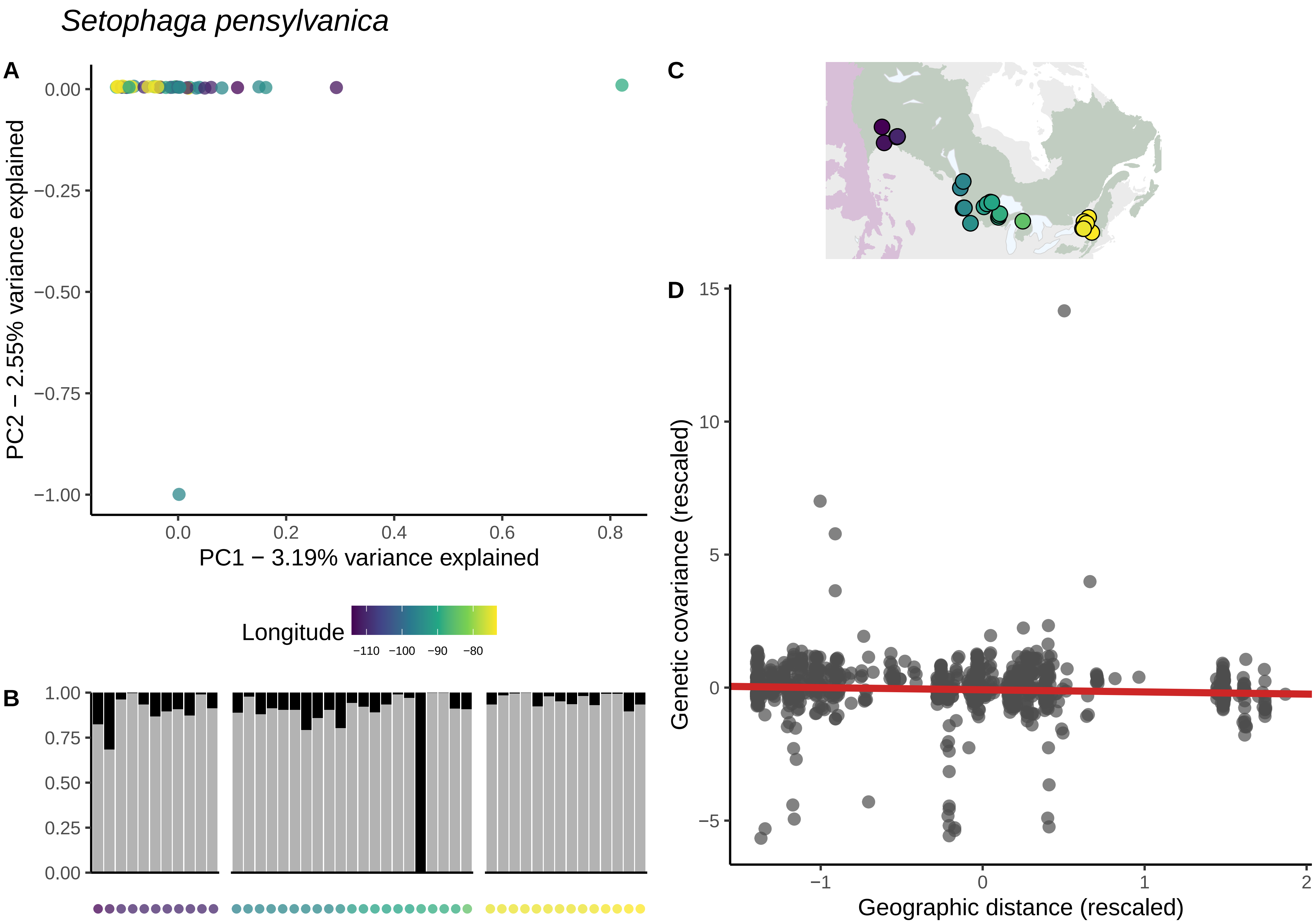

Setophaga pensylvanica
A
PC2 − 2.55% variance explained
PC1 − 3.19% variance explained
0.00
−0.25
−0.50
−0.75
−1.00
0.0
0.2
0.4
0.6
0.8
Longitude
−110
−100
−90
−80
B
1.00
0.75
0.50
0.25
0.00
C
D
Genetic covariance (rescaled)
Geographic distance (rescaled)
15
10
5
0
−5
−1
0
1
2

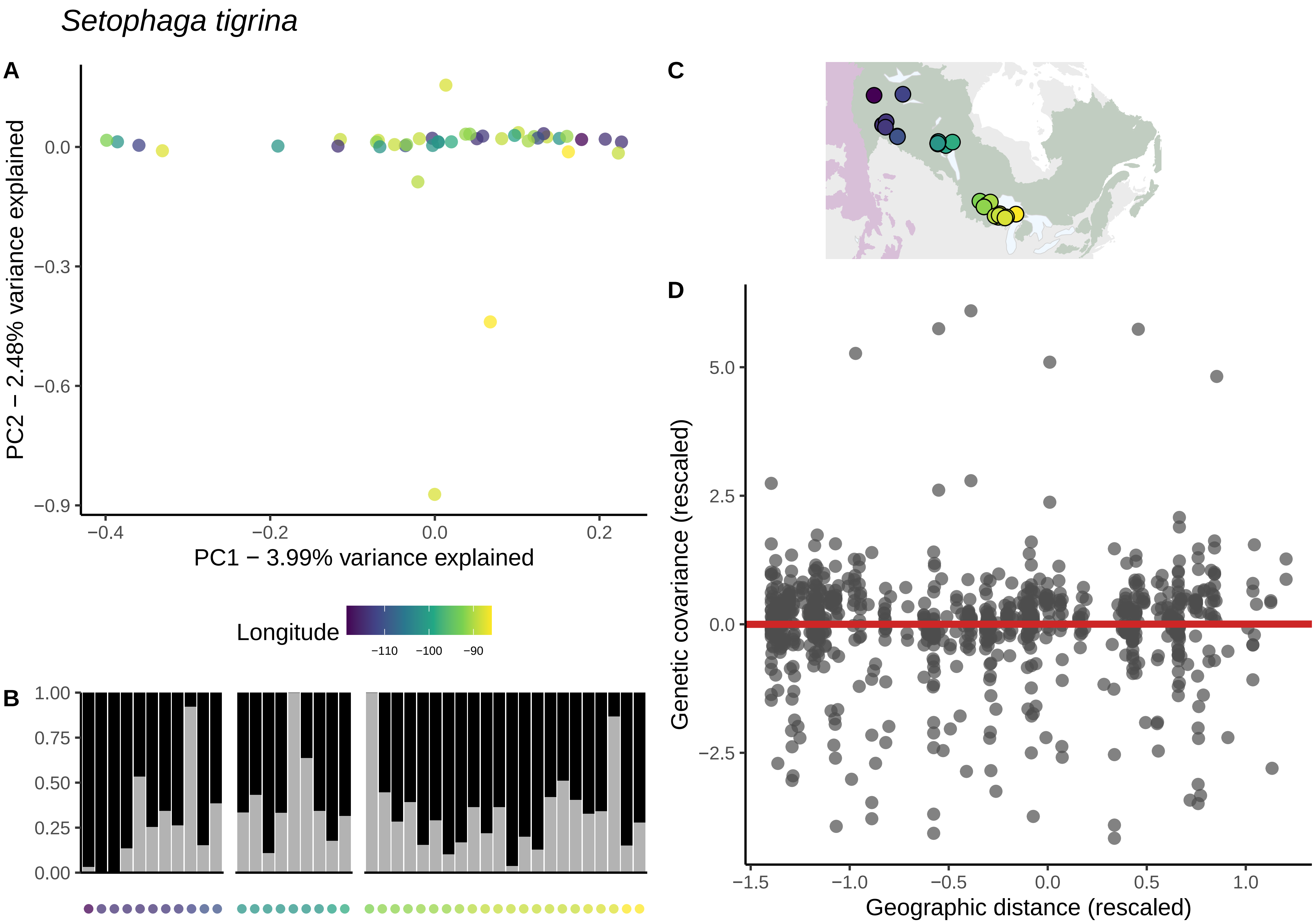

Setophaga tigrina
A
PC2 − 2.48% variance explained
PC1 − 3.99% variance explained
Longitude
−110
−100
−90
B
C
D
Genetic covariance (rescaled)
Geographic distance (rescaled)

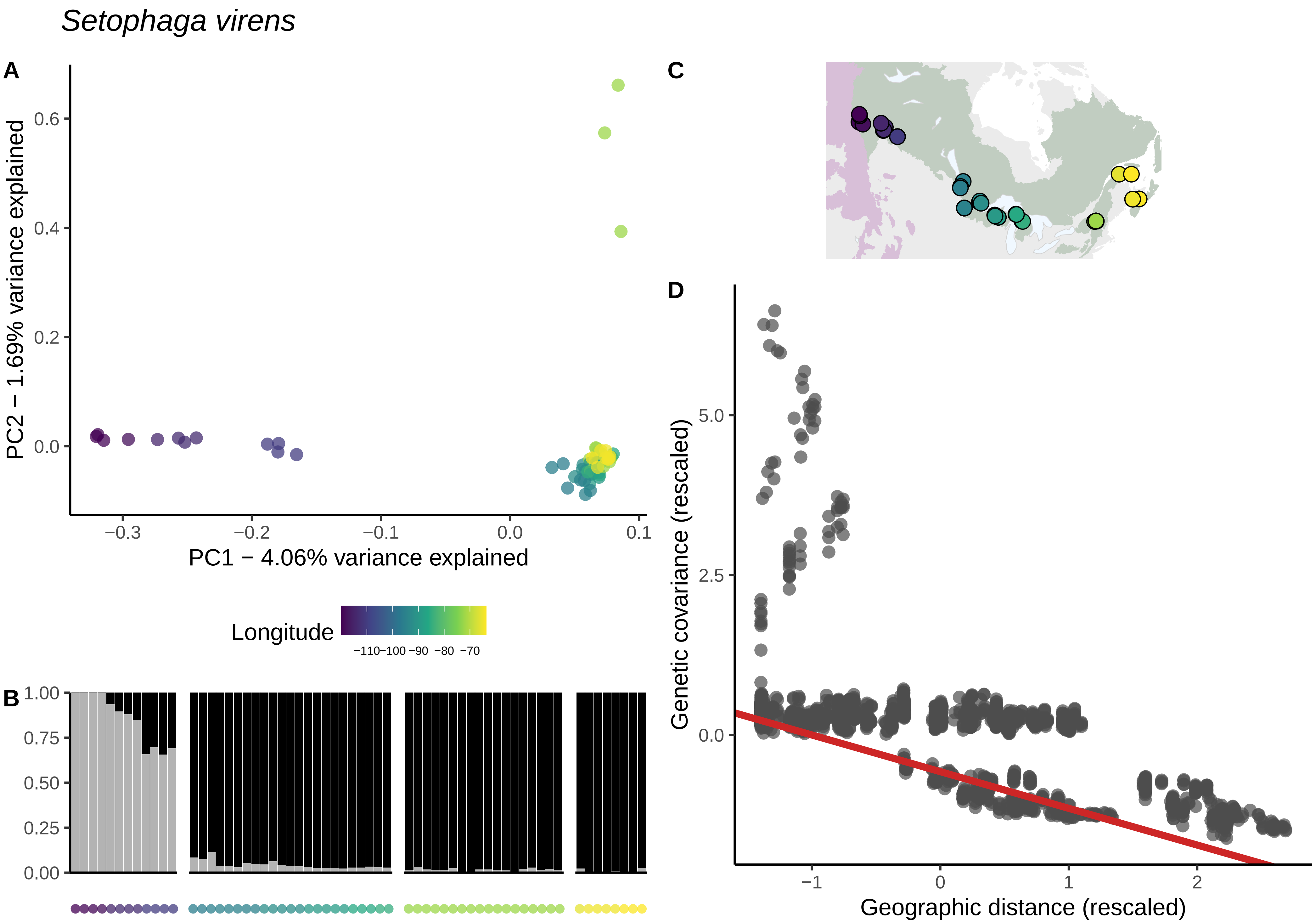

Setophaga virens
A
PC2 − 1.69% variance explained
0.6
0.4
0.2
0.0
−0.3
−0.2
−0.1
0.0
0.1
PC1 − 4.06% variance explained
Longitude
−110−100 −90 −80 −70
B
1.00
0.75
0.50
0.25
0.00
C
D
Genetic covariance (rescaled)
5.0
2.5
0.0
−1
0
1
2
Geographic distance (rescaled)
